# An ncBAF-ETS2 Chromatin-Remodelling Axis Drives Vascular Smooth Muscle Cell Osteogenic Reprogramming in Vascular Calcification

**DOI:** 10.64898/2026.08.21.746239

**Authors:** Meng-Ying Wu, Jirapath Thammaphet, Audrey Kelly, Samara Banday, Sadia Ahmad, Chin-Yee Ho, Sujin Lee, Elizabeth Moore, Rajeev Malhotra, Clint L. Miller, Konstantinos Theofilatos, Paul Lavender, Andrew Durham, Catherine M. Shanahan

## Abstract

**Introduction:** Vascular calcification is a detrimental ageing-related pathology that is markedly accelerated in metabolic disorders. It is driven by osteogenic differentiation of vascular smooth muscle cells (VSMCs), however epigenetic regulatory pathways activated early in this transition remain poorly defined.

**Methods:** An in vitro calcification model was developed using primary human aortic VSMCs cultured with or without mineral stress. Epigenetic changes were assessed using targeted PCR arrays and CUT&RUN sequencing. Key findings were validated in vivo using single-cell sequencing datasets from human large arteries and spatial transcriptomic analysis in atherosclerotic carotid plaques. Transcriptomic and CUT&RUN analyses identified gene targets altered by epigenetic remodelling, and molecular tools were applied to study effects on metabolism, inflammation, apoptosis, and calcification.

**Results:** During early calcification in response to mineral stress, SWI/SNF chromatin remodelling complexes shift toward ncBAF enrichment in pre-osteogenic VSMCs. ncBAF complexes activated transcriptional programs involved in inflammation, apoptosis, and glycolysis—all hallmarks of calcifying VSMCs. The transcription factor ETS2 was identified as a novel component of ncBAF complexes. Disruption of ncBAF or ETS2 impaired osteogenic differentiation and calcification. Notably, ETS2 expression was regulated by ncBAF, forming a positive feedback loop that reinforced VSMC phenotypic switching. Co-activation of ETS2 and ncBAF and the resulting transcriptional shifts were confirmed in human arterial single-cell datasets, with osteogenic/inflammatory clusters showing NFkB and RUNX2 activation. Spatial transcriptomics further suggested that a macrophage-rich microenvironment may promote the differentiation of smooth muscle cells toward an overt osteogenic/inflammatory phenotype. Immunohistochemistry showed that ETS2 levels correlated with calcification severity in human vessels supporting the potential clinical relevance of ETS2.

**Conclusions:** Our findings identify a novel epigenetic mechanism in vascular calcification, where ncBAF and ETS2 cooperate to drive VSMC phenotypic switching. This ncBAF–ETS2 axis represents a potential therapeutic target to modulate VSMC plasticity and intervene early in the progression of cardiovascular calcification.

## Introduction

Vascular calcification (VC) is strongly linked to an increased risk of cardiovascular mortality^1^. It is prevalent in the vessel intima in atherosclerosis and markedly accelerated in the vessel media in chronic kidney disease (CKD) where it is driven by dysregulated mineral metabolism^2^. VC is a cell-mediated process associated with vascular smooth muscle cell (VSMC) apoptosis, phenotypic modulation and maladaptation^3,4^. VSMC contractile activity is essential for the optimal function of blood vessels, primarily maintaining blood pressure^5^ and VSMCs also play a vital role in maintaining and remodelling the extracellular matrix (ECM)^6^. The plasticity of VSMCs allows them to dedifferentiate in response to environmental cues. In calcified vessels, characterized by the deposition of apatite mineral in the arterial wall, VSMCs exhibit osteo/chondrogenic differentiation, which is triggered by transcriptional changes that repress contractile markers and increase expression of osteogenic factors usually expressed during bone formation^7–9^. Single-cell meta-analysis of human atherosclerotic artery samples have uncovered the complexity of VSMC phenotypic changes and pseudotime trajectory analysis has demonstrated that osteogenic markers are elevated at the end-stage of atherosclerosis progression^10^. However, little is known of the epigenetic changes that precede the onset of the transcriptional changes that drive VSMC dedifferentiation.

SWI/SNF chromatin remodelling complexes comprise an evolutionarily conserved family of ATP-dependent remodellers that hydrolyse ATP to modulate DNA–protein interactions^11^, thereby regulating access of the transcriptional machinery to chromatin and controlling gene expression ^12,13^. Three major SWI/SNF subtypes—canonical BAF, PBAF, and non-canonical BAF (ncBAF)—are defined by their unique subunit compositions. BAF complexes are characterized by the mutually exclusive incorporation of ARID1A or ARID1B, members of the ARID protein family containing a conserved AT-rich interaction domain^14^. PBAF complexes are distinguished by ARID2, polybromo-1 (PBRM1), and bromodomain-containing protein 7 (BRD7), with PBRM1 and BRD7 exhibiting high affinity for acetylated lysine residues^15^, including the active chromatin mark histone H3 lysine 27 acetylation (H3K27ac)^16,17^. In contrast, ncBAF complexes are defined by the presence of bromodomain-containing protein 9 (BRD9) which also has high affinity for H3K27ac^18^.

SWI/SNF complexes are essential regulators of pluripotency, cellular reprogramming, and lineage specification ^19–22^, and are well established as tumour suppressors across diverse human cancers ^23,24^. Emerging evidence further implicates SWI/SNF components in the regulation of VSMC plasticity. The shared catalytic subunit BRG1 has been shown to cooperate with histone deacetylase HDAC9 to repress contractile gene expression in VSMCs ^25^, whereas the core subunit BAF60A is upregulated in human abdominal aortic aneurysm lesions, where it promotes inflammatory activation and ECM degradation ^26^. Because disruption of transcriptional regulation broadly drives VSMC dedifferentiation, we hypothesized that SWI/SNF chromatin-remodelling complexes are required for VSMC osteogenic differentiation. We further hypothesized that distinct SWI/SNF complex subtypes exert non-redundant regulatory functions during VSMC dedifferentiation.

To address these questions, we investigated SWI/SNF complex composition across the timecourse of calcification. We show that SWI/SNF complexes undergo subunit switching during VSMC osteogenic processes and that ncBAF in cooperation with ETS2 promotes osteogenic priming of VSMCs with both co-activated in pre-osteogenic and osteogenic/inflammatory VSMC subpopulations occupying distinct localisations in atherosclerotic plaques. These results identify a novel ETS2–ncBAF axis as a key regulator of VSMC plasticity and a potential therapeutic target in early vascular calcification.

## Results

### The ncBAF unique component BRD9 and the catalytic subunit BRG1 mediate osteogenic differentiation of VSMCs

To study the impact of epigenetic mechanisms on osteogenic differentiation in VSMCs, we performed targeted transcriptome analysis by QPCR array, to screen expression levels of 160 epigenetic regulators across the time course of mineral stress induced calcification (Fig.S1A-B). VSMCs were exposed to osteogenic media, and the calcification stages were defined as previously^27,28^ described: pre-stage was 48 hours after exposure to osteogenic media (no mineralization), early-stage was when visible microcalcification on alizarin red staining was observed and late-stage corresponded to extensive deposition of mineral. Calcium quantification showed stepwise increases as calcification progressed (Fig.S1B).

Epigenetic regulator screening showed that a panel of genes belonging to the Bromodomain (BRD) protein family were upregulated during calcification progression (Fig.1A). Compared to control VSMCs in normal media, *BRD2* and *BRD3* expression rapidly increased at the pre-stage and continued to increase as calcification progressed. *BRD1*, *BRD4*, and *BRD8* were downregulated at the pre-stage but then increased in early and late stage calcification. In addition, a set of components of the SWI/SNF chromatin remodelling complexes showed dramatic changes during VSMC calcification (Fig.1A). Two catalytic subunits, *BRG1* and *BRM*, that are mutually exclusive within SWI/SNF complexes, showed differential expression; *BRM* was repressed throughout calcification progression while *BRG1* was induced in late-stage calcification. Moreover, two subtypes of the SWI/SNF complex, PBAF and BAF, displayed distinct expression patterns in calcified VSMCs. *BRD7* and *PBRM1*, two unique subunits of the PBAF complex, were both upregulated in late-stage calcification. In contrast, *ARID1A*, a defined subunit of BAF complexes, was upregulated in pre-stage calcification, but repressed at later stages suggesting the possibility that Arid1a was replaced from BAF complexes as calcification progressed. Taken together, these data implicate BRD proteins and SWI/SNF chromatin remodelling complexes in VSMC phenotypic modulation in response to calcification stimuli.

**Figure 1.**
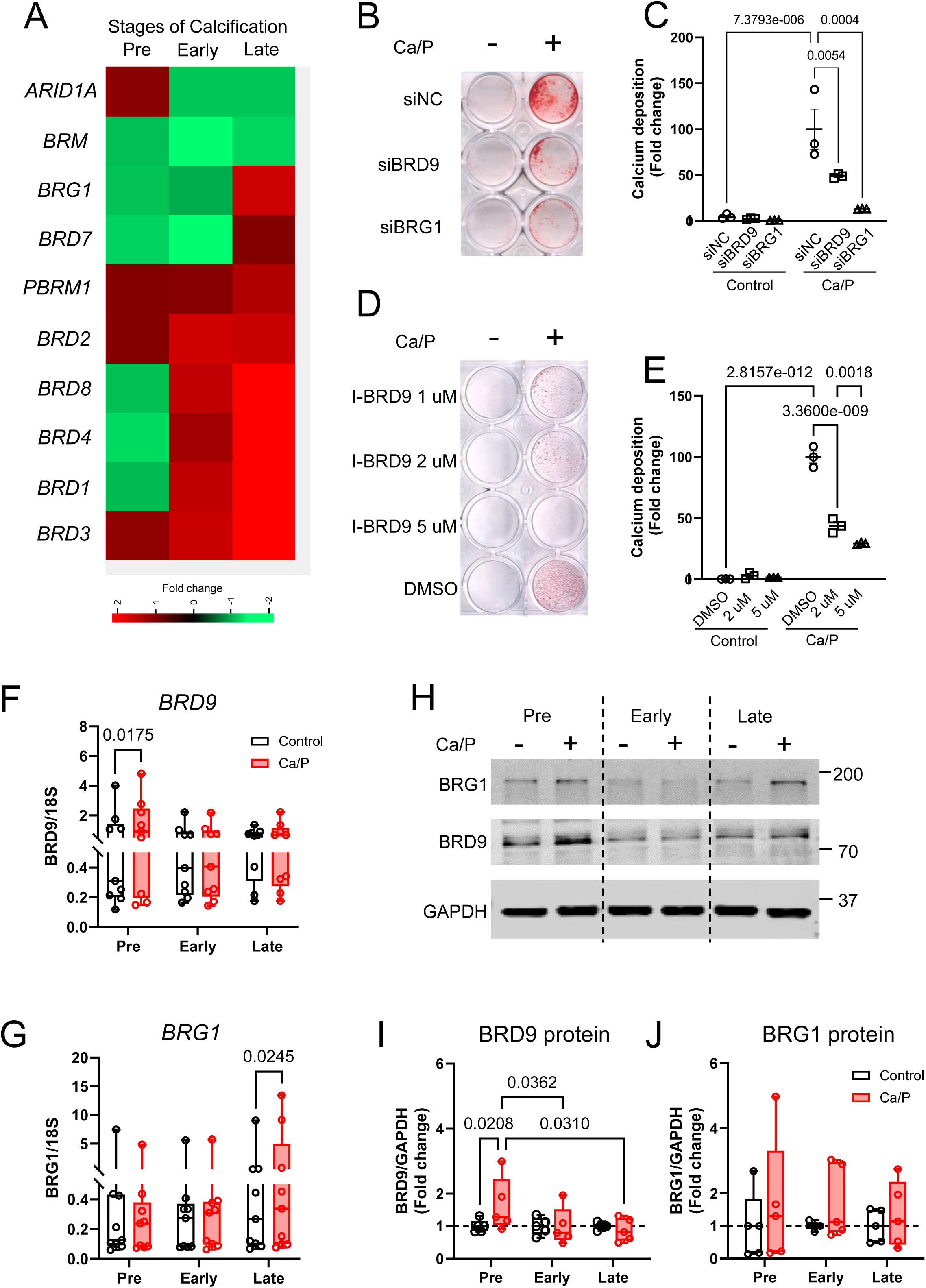
ncBAF complex components BRD9 and BRG1 function to promote osteogenic differentiation in human vascular smooth muscle cells (VSMCs). (A) Heatmap showing gene expression profiles of epigenetic regulators during calcification progression. (B) Alizarin red staining at late-stage calcification in VSMCs after depletion of BRD9 and BRG1 and treatment with control or osteogenic media with elevated calcium and phosphate (Ca/P) to induce calcification. (C) o-Cresolphthalein assay was used to quantify calcium content of VSMCs at late-stage calcification after BRD9 and BRG1 depletion. Total calcium content was normalized to total protein content in each sample. N= 3 independent experiments from 3 isolates. Significance was analyzed by Mixed-effects model test and q-values are shown. (D) Alizarin red staining at late-stage calcification of VSMCs cultured with control or osteogenic media with different concentrations of the BRD9 inhibitor (I-BRD9). (E) o-Cresolphthalein assay was used to quantify calcium content of VSMCs at late-stage of calcification after culture with control or osteogenic media with different concentrations of the BRD9 inhibitor (I-BRD9). N= 3 independent experiments from 3 isolates. Significance was analyzed by Mixed-effects model test and q-values are shown. (F-G) Detection for *BRD9* and *BRG1* expression in control or calcified VSMCs at the 3 stages of calcification by QPCR. N= 9 independent experiments from 3 isolates. Significance was analyzed by Mixed-effects model test and q-values are shown. (H) Western blotting showing protein expression profiles of BRD9 and BRG1 in control or calcified VSMCs at the 3 stages of calcification. (I-J) Quantification of the protein levels of BRD9 and BRG1 in control or calcified VSMCs at 3 stages of calcification. N= 5 independent experiments from 3 isolates. Significance was analyzed by Mixed-effects model test and q-value are shown.

To determine the role of individual BRD proteins during VSMC calcification, we systematically depleted each BRD protein using siRNA treatment. This showed that only depletion of BRD9 significantly attenuated VSMC mineralization (Fig. S1C). As BRD9 is a defined component of ncBAF (non-canonical BAF) complexes^18^, we subsequently knocked-down a set of unique components belonging to different subtypes of SWI/SNF complexes, as well as the common catalytic subunit BRG1 (Fig.S1D-H). We found that depletion of BRG1 and BRD9 had a dramatic effect on blocking VSMC calcification (Fig. 1B-C). In contrast, depleting PBRM1 and BRD7 showed an enhancement of calcification, compared to cells treated with scramble siRNA negative control (Fig S1D). Knock down of ARID1A also blocked calcification but was less effective than BRG1 and BRD9 depletion (Fig S1D, Fig1B-C) suggesting that BRD9 and BRG1 within ncBAF complexes function predominately on VSMC mineralization.

To confirm the role of BRD9 in calcification we next used a chemical probe I-BRD9 to block BRD9 binding affinity to acetylated histone marks. I-BRD9 dramatically reduced VSMC calcification in a dose-dependent manner (Fig. 1D-E), suggesting BRD9 promotes calcification via its association with acetylated histone marks on the chromatin landscape^29^. As *BRD9* was not included in the targeted transcriptomic QPCR array, we next monitored *BRD9* expression throughout VSMC calcification. *BRD9* was upregulated in pre-stage calcification (Fig. 1F), indicating *BRD9* responds early in the calcification cascade. We also confirmed *BRG1* expression was upregulated in late-stage calcification (Fig. 1G), consistent with QPCR array data. Immunoblotting showed that BRD9 protein increased at the pre-stage of calcification, followed by decreasing expression during calcification progression (Fig. 1H-I). In contrast, the protein level of BRG1 showed no dramatic changes between calcifying and control VSMCs (Fig. 1J). These data suggest that BRG1 is stably expressed in both contractile and calcifying VSMCs while BRD9 is an early marker of the initiation of VSMC phenotypic change and may indicate an increased abundance of ncBAF complexes in pre-osteogenic VSMCs.

### ncBAF complexes regulate the expression of osteogenic and inflammatory markers in calcified vascular smooth muscle cells

To determine the mechanism whereby BRD9 functions to regulate VSMC calcification, we examined the effect of inhibiting BRD9 on expression of osteogenic markers during the progression of mineralization. I-BRD9 treatment of VSMCs reduced osteogenic markers *BMP2*, *MSX2* and *RUNX2* (Fig 2A) and depleting BRD9 mirrored the effect of I-BRD9 treatment (Fig S2A). Loss of BRG1 showed a similar effect to depleting BRD9, reducing the expression of *BMP2*, *MSX2*, *RUNX2* (Fig 2B). The contractile marker *TAGLN* was decreased in late-stage calcification (Fig S2B) and I-BRD9 repressed *TAGLN* expression in both control and calcifying conditions (Fig S2B) while depleting BRD9 or BRG1 showed no effect (Fig S2B). We also examined the effects of functional blocking of BRD9 on inflammatory factors previously shown to be linked to VSMC osteogenic differentiation^27,30,31^. Interleukin 6 (*IL6*) expression increased in VSMCs from early to late-stage calcification (Fig 2C), and inhibiting BRD9 or BRG1 efficiently reduced *IL6* in late-stage calcification (Fig 2C).

**Figure 2.**
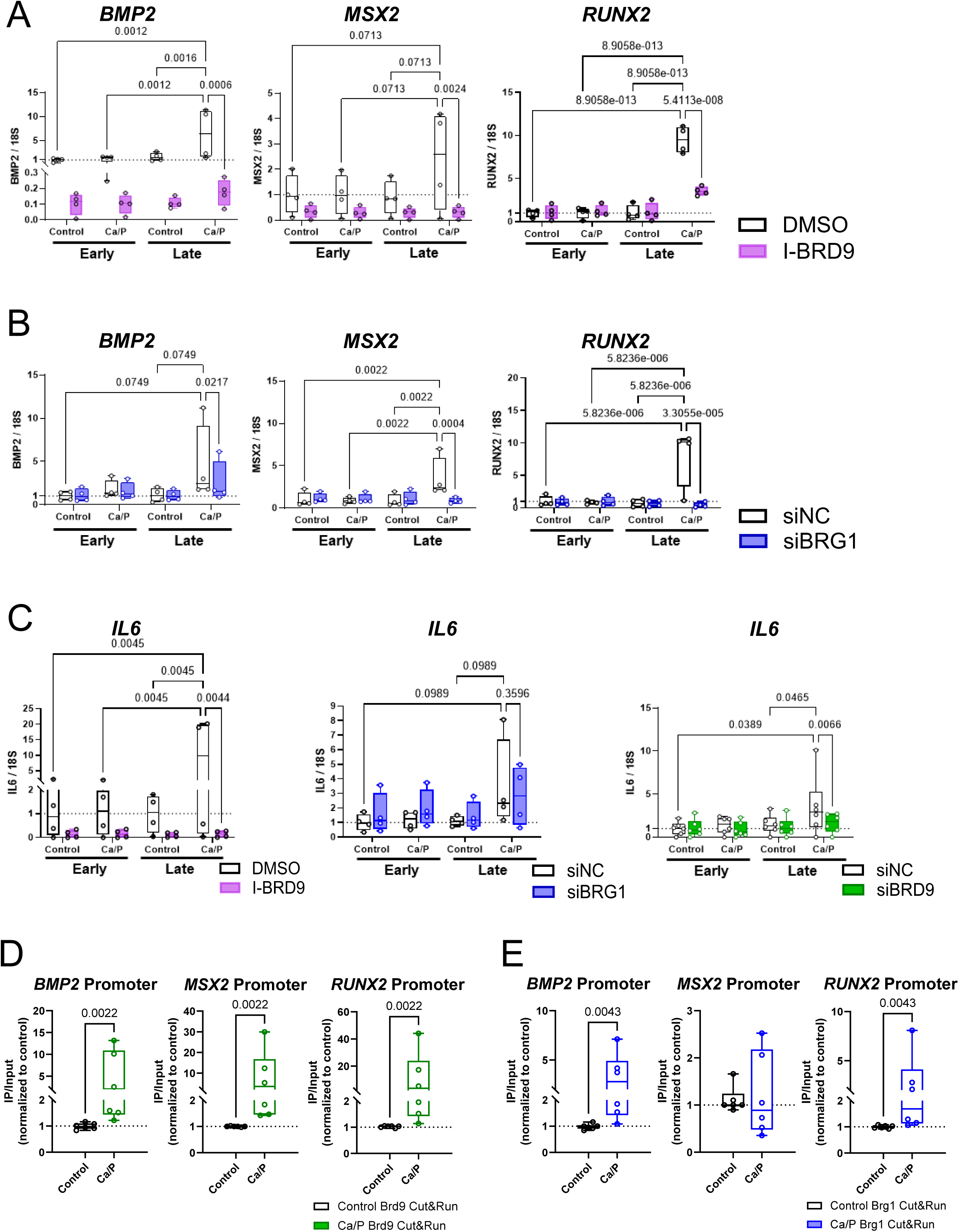
Epigenetic influences by ncBAF complex components Brd9 and Brg1 regulated osteogenic and inflammatory markers in VSMCs under calcification stimuli. (A) I-BRD9 treatment repressed *BMP2*, *MSX2*, *RUNX2* expression at late-stage calcification in VSMCs. VSMCs were cultured in control or osteogenic media with the presence or absence of I-BRD9. Samples were collected at the early- or late-stage of calcification. N= 4 independent experiments from 2 isolates. Significance was analyzed by Mixed-effects model test and q-values are shown. (B) Loss of BRG1 by RNAi treatment repressed *BMP2*, *MSX2*, *RUNX2* expression at late-stage calcification in VSMCs. VSMCs were transfected by non-targeting siRNA (siNC) or BRG1-targeting siRNA (siBRG1) under osteogenic stimuli. Samples were collected at the early- or late-stage of calcification. N= 4 independent experiments from 2 isolates. Significance was analyzed by Mixed-effects model test and q-values are shown. (C) Repression of *IL6* expression by inhibiting BRD9 and BRG1 at late-stage calcification in VSMCs. N= 4 independent experiments from 2 isolates for experiments with I-BRD9 treatment and loss of BRG1 by RNAi. N= 6 independent experiments from 2 isolates for experiment with depletion of BRD9 by RNAi. Significance was analyzed by Mixed-effects model test and q-values are shown. (D-E) The enrichment of BRD9 (D) and BRG1 (E) at the promoters of *BMP2*, *MSX2*, and *RUNX2* in VSMCs under osteogenic stimuli detected by ChIP-QPCR. N= 6 independent experiments from 2 isolates. Significance was analyzed by Mann-Whitney test and p-value are shown.

Next we tested whether BRD9 and BRG1 directly mediate the expression of osteogenic genes and performed Cleavage Under Targets and Release Using Nuclease (Cut&Run) followed by RT-qPCR which revealed that the occupancy of BRD9 on promoters of *BMP2*, *MSX2* and *RUNX2* was increased in VSMCs under osteogenic stress compared to control (Fig 2D). Increased occupancy of BRG1 at promoters of *BMP2* and *RUNX2* was also observed (Fig 2E) suggesting that BRD9 and BRG1 as part of ncBAF complexes are involved in osteogenic differentiation of VSMCs by directly regulating the activation of osteogenic markers with little or no role in contractile gene expression.

### Subunit switching within SWI/SNF chromatin remodelling complexes and novel interactors define distinct VSMC phenotypes

We next set out to identify the composition of SWI/SNF complexes during VSMC osteogenic phenotypic modulation using Immunoprecipitation (IP) followed by mass spectrometry analysis (IP-MS)^32,33^. In control VSMCs, BRG1-IP co-purified with most components of SWI/SNF complexes (Fig 3A), including the core components SNF5, BAF170, BAF57, BAF53A, BAF60B/C, BCL7C, and unique subunits, such as ARID1A/B (representative of BAF), PBRM1 (representative of PBAF), and BRD9 (representative of ncBAF). In VSMCs treated with osteogenic media for 5 days (Fig 3A Ca/P) some core components were absent from the BRG1-IP; only BAF170, BCL7C, and BAF53A together with an additional core subunit BAF155 were pulled down. Moreover, the unique subunits ARID1A/B were reduced from the BRG1-IP, suggesting that loss of ARID1A/B within SWI/SNF complexes occurs when VSMCs transform into a pre-osteogenic phenotype. Meanwhile, BRD9 and BRD7 were enhanced in VSMCs under osteogenic stimuli suggesting subunit switching within SWI/SNF complexes is a feature of VSMC early osteogenic change.

**Figure 3.**
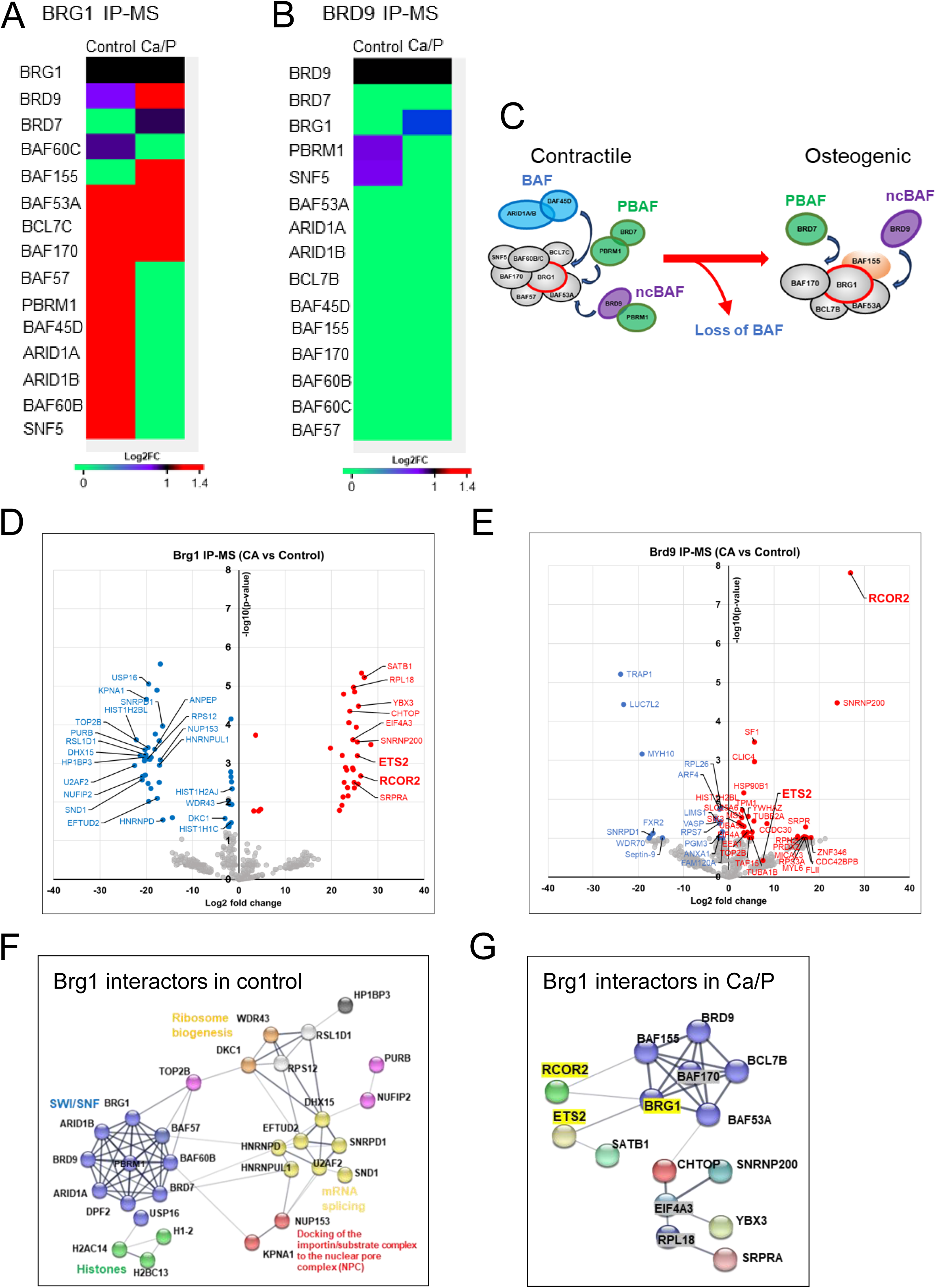
Compositional changes in the subtypes of SWI/SNF complexes and specific interactors in distinct phenotypes of VSMCs defined by IP-mass spectrometry analysis. (A-B) Changes to the composition of SWI/SNF complexes in VSMCs cultured with control or osteogenic media (Ca/P) for 5 days. The heatmaps indicate the log2 fold changes of protein abundance of SWI/SNF subunits. All values are the average of 2 repeats from 2 different isolates. Protein intensities were normalized to the bait protein to present as stoichiometric interactions for studying composition changes across different conditions. (C) Colour scheme showing subunit switching within SWI/SNF complexes during osteogenic differentiation. Three subtypes of SWI/SNF complexes were exhibited in control VSMCs. Reducing BAF complexes and increasing PBAF and ncBAF complexes were found in pre-osteogenic VSMCs. (D-E) Volcano plots show proteins enriched in BRG1 IP-MS (D) and BRD9 IP-MS (E) in VSMCs with presence or absence of osteogenic stimuli. Using two sample t-test to analyse the statistically significant differences in protein intensities in IP samples in normal VSMCs or pre-osteogenic VSMCs and volcano plots (cutoff at Log2FC > 4 and – log_10_(p-value) > 1.3) to show novel interactors of BRG1 or BRD9 in pre-osteogenic VSMCs (red dots) or Control (blue dots). (F-G) String network demonstrated interactors of BRG1 in control VSMCs (F) and pre-osteogenic VSMCs (G).

As BRD9 and BRD7 both bound to BRG1 in pre-osteogenic VSMCs, we next performed BRD9 IP-MS to determine what complexes were present. BRD9-IP co-purified with BRG1 in pre-osteogenic VSMCs but not in control, suggestive of an enhanced interaction between BRD9 and BRG1 during early osteogenic change (Fig 3B). BRD7 was absent in the BRD9-IP in both normal and osteogenic VSMCs (Fig 3B) indicating BRD7 and BRD9 associate with different complexes consistent with findings in other cell types^18^. Additionally, we unexpectedly found PBRM1 interacted with BRD9 in normal VSMCs (Fig 3B), which does not concur with the current notion of the composition of PBAF complexes^18^ suggesting this may be a VSMC specific interaction.

IP using BRG1 or BRD9 antibodies followed by Western blotting (IP-WB) was used to confirm these interactions. Reciprocal pull-down of each component in VSMCs cultured in both control or osteogenic media indicated that ncBAF complexes comprised of BRG1 and BRD9 were present in both conditions (Fig S3A). We also confirmed the interaction between BRD9 and PBRM1 in both control and osteogenic VSMCs (Fig S3B).

Taken together these data suggest that the known SWI/SNF subtypes (BAF, PBAF, ncBAF) exist in normal contractile VSMCs (Fig 3C). During early VSMC osteogenic change the subunits switch leading to a loss of BAF (defined by ARID1A/B) but maintenance of both ncBAF and PBAF complexes (Fig 3C).

In addition to the above, BRG1 IP-MS analysis also identified a group of novel interactors of BRG1 in normal VSMCs and in response to osteogenic stimuli. Volcano plot analysis showed 45 proteins were significantly enriched in BRG1-IP in control VSMCs (Fig 3D, blue dots). 22 of these proteins were identified as nuclear proteins which possibly associate with SWI/SNF to function as regulators of chromatin structure. STRING network analysis^34^ revealed that they contribute to diverse cellular functions, such as histone deubiqutinase USP16, histone H2A, H2B, H1.2 and DNA topoisomerase TOP2B which functions to regulate chromatin structure. SWI/SNF complexes also associated with the nuclear pore complex (NPC) subunit NUP153 and NPC docking substrate KPNA1, which function as molecular traffic regulators. Proteins involved in mRNA splicing and ribosome biogenesis were also found to interact with SWI/SNF complexes in normal VSMCs (Fig 3F).

We also identified 33 proteins significantly enriched in BRG1-IP in VSMCs under osteogenic stimuli (Fig 3D, red dots) and 9 of these proteins were identified as nuclear proteins. STRING network analysis showed that these 9 proteins have been reported to interact with SWI/SNF complexes through subunits BRG1, BAF155, or BAF53A (Fig 3G). Among these 9 proteins, RCOR2 and ETS2 also strongly interacted with BRD9 by IP-MS in pre-osteogenic VSMCs (Fig 3E), suggesting that they are novel components of ncBAF complexes specifically in pre-osteogenic VSMCs. RCOR2 has been reported to work with histone demethylase LSD1, which regulates the levels of histone H3K4 or K9 mono- or di-methylation and shows dual functions as an activator or repressor^35,36^, suggesting ncBAF function may be related to the level of histone methylation regulated by RCOR2-LSD1 complexes. ETS2 functions as a transcription factor, and pro-oncogene^37,38^ and regulates genes involved in development, apoptosis, and inflammatory responses^39,40^. These interactors suggest that ncBAF complexes are specifically involved in several biological functions including programmed cell death and inflammatory activation under osteogenic conditions.

### Transcriptional regulation of ncBAF complexes leads to changes in the molecular signature of pre-osteogenic VSMCs

We next explored the active genomic regions directly regulated by ncBAF complexes during VSMC osteogenic change by subjecting VSMCs cultured in control or osteogenic media for 5 days to Cut&Run-Seq analysis^41^ with antibodies against endogenous BRD9 and BRG1, as well as the active histone mark H3K27ac. Peak annotation in DiffBind analysis showed that nearly 50% and 20% of enriched peaks of BRD9 and BRG1 respectively, in pre-osteogenic VSMCs, were in promoter regions (Fig S4A), indicating BRD9 and BRG1 are likely to directly remodel transcriptional patterns in VSMCs under osteogenic stimuli. BRD9-enriched regions in pre-osteogenic VSMCs showed greatly reduced BRD9 binding intensity in control VSMCs (Fig S4B). BRG1 localization was also changed between control and pre-osteogenic VSMCs (Fig S4B) suggesting distinct localization of ncBAF complexes in association with VSMC phenotype switching.

We next overlayed peaks of co-localization for BRD9, BRG1, and H3K27ac and identified 250 activated genes that are potentially regulated by ncBAF complexes in pre-osteogenic VSMCs (Fig 4A). MSiDB Hallmark analysis on Enrichr software^42,43^ revealed these 250 genes were strongly related to TNFα signalling via NFkB activation, as well as reactive oxygen species pathway, KRAS signalling and response to hypoxia (Fig 4B). To determine whether ncBAF targeted genes play a role in VSMC mineralization, we compared the CUT&RUN-Seq analysis to RNA-seq data for control and pre-osteogenic VSMCs. 1536 differentially expressed genes were defined by RNA-seq analysis (log2FC=0.5 and -log10 (p-value) > 0.7), of which1017 and 519 genes were upregulated and downregulated, respectively, in VSMCs under osteogenic stimuli. MSiDB Hallmark analysis showed upregulated genes were related to TNFα signalling via NFkB activation, epithelial mesenchymal transition, response to hypoxia and inflammatory responses (Fig 4D), and down-regulated genes were involved in UV response, epithelial mesenchymal transition, and the p53 pathway (Fig 4E). 23 of these 1017 up-regulated genes showed differential peaks in the ncBAF CUT&RUN-Seq (Fig 4F). Interestingly, these 23 genes were also highly related to TNFα signalling via NFkB activation (Fig 4G), the most up-regulated pathway in calcified VSMCs by RNA-seq. Importantly, one of the ncBAF targeted genes was *ETS2*, which was also identified as a novel interactor with ncBAF (Fig 3D-E). We validated whether the expression of *ETS2*, as well as five additional genes involved in the NFkB activation pathway, including *TNFAIP3*, *CXCL1*, *PTGS2*, *BCL2A1* and *PFKFB3* (Fig S4C) were regulated by BRD9 by treating VSMCs with I-BRD9 in osteogenic media. *TNFAIP3*, *ETS2*, *CXCL1*, *PTGS2*, and *BCL2A1* were up-regulated in late-stage calcification and were repressed by I-BRD9 treatment (Fig 4H-L). *PFKFB3* showed no change in late-stage calcification, however, it was also significantly repressed by I-BRD9 (Fig 5M). Notably, in addition to NFkB activation ^30,31,44–46^ these same 5 genes have been implicated in additional functions relevant to vascular calcification. *BCL2A1*, *PTGS2*, and *ETS2* have been shown to play a role in regulating apoptosis ^39,47,48,49^ while *PFKFB3*, encodes a key metabolic enzyme in glycolysis ^50^ suggesting ncBAF complexes remodel transcriptional patterns to activate diverse molecular functions in VSMCs relevant to osteogenic processes.

**Figure 4.**
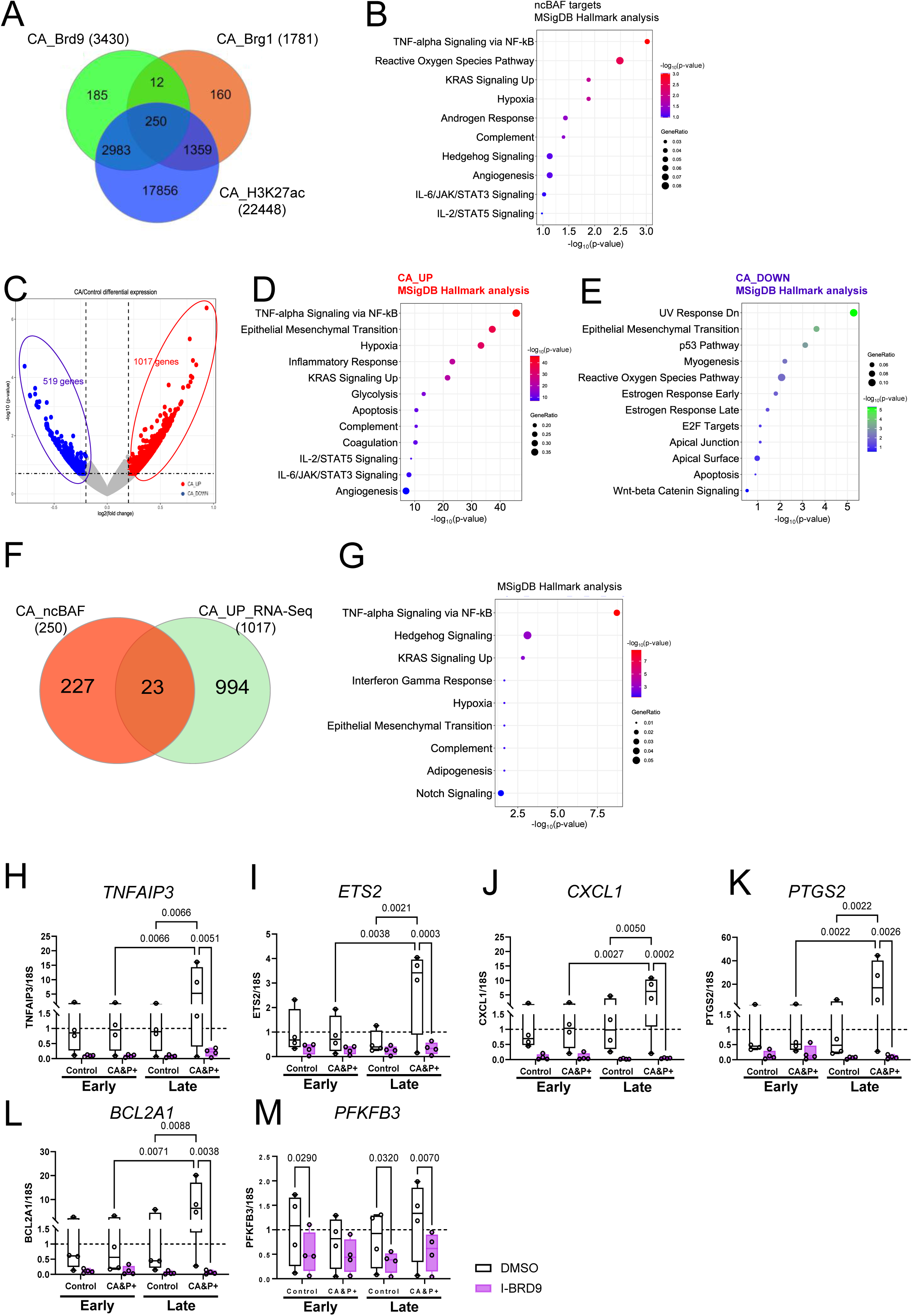
ncBAF Complexes remodelled the transcriptional signatures in VSMCs under osteogenic stimuli. (A) Genome-wide BRD9 targets (3430 genes), BRG1 targets (1781 genes), and active mark H3K27ac locations were analysed by Cut&Run-Seq in VSMCs in response to osteogenic stimuli. The intersection of these three datasets showed 250 active ncBAF-dependent genes in pre-osteogenic VSMCs. (B) MSigDB Hallmark analysis of 250 ncBAF-dependent genes revealed associations with TNFα signalling via NFkB activation, Reactive Oxygen Species pathway, and KRAS Signalling Up as top scoring terms, respectively. (C) A volcano plot shows significantly upregulated and downregulated genes in pre-osteogenic VSMCs compared to control. Cut off: log2FC=0.5 with p-value=0.05. (D) MSigDB Hallmark analysis of upregulated genes in pre-osteogenic VSMCs showed associations with TNFα signalling via NFkB activation, Epithelial Mesenchymal Transition, and Hypoxia as top scoring terms, respectively. (E) MSigDB Hallmark analysis of downregulated genes in pre-osteogenic VSMCs showed associations with UV response, Epithelial Mesenchymal Transition, and p53 pathway as top scoring terms, respectively. (F) Overlap between ncBAF-dependent genes (250) and upregulated genes (1017) in pro-osteogenic VSMCs confirmed 23 ncBAF-dependent genes were upregulated in VSMCs response to osteogenic stimuli. (G) MSigDB Hallmark analysis of 23 genes in pre-osteogenic VSMCs showed associations with TNFα signalling via NFkB activation, Hedgehog Signalling, and KRAS Signalling Up as top scoring terms, respectively. (H-M) Treatment with the BRD9 inhibitor (I-BRD9) repressed the expression of 6 ncBAF-dependent genes involved in NFkB activation validated by QPCR. N= 4 independent experiments from 2 isolates. Significance was analyzed by Mixed-effects model test and q-values are shown.

**Figure 5.**
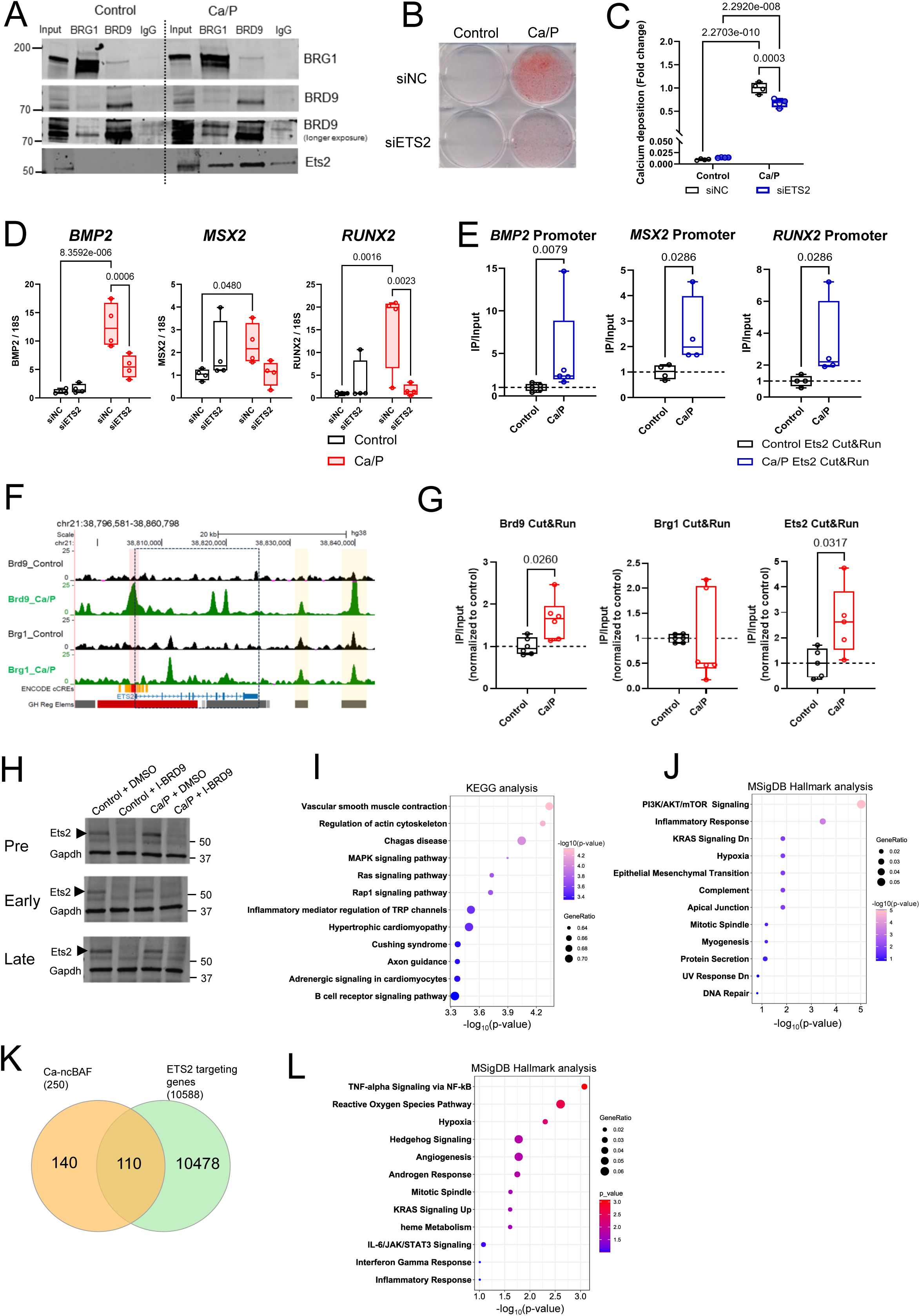
ETS2 expression requires the function of ncBAF complexes and is involved in activating different pathways in VSMCs under osteogenic stimuli. (A) ETS2 interacting with ncBAF (BRG1-BRD9) complexes in pre-osteogenic VSMCs was confirmed by immunoprecipitation (IP) followed by western blotting. Whole cell lysates were made from VSMCs cultured in control or osteogenic media for 5 days. Antibodies against BRG1 or BRD9 were pre-bound with Protein A-dynabeads before incubation with 750 µg of whole cell extract. IP using Rabbit IgG was applied as a negative control. Capture of BRG1, BRD9 and ETS2 was confirmed by western blotting. (B-C) Depleting ETS2 by RNAi treatment attenuated osteogenic differentiation in VSMCs demonstrated by Alizarin Red staining (B) and (C) o-Cresolphatalein assay. (D) ETS2 regulated the gene expression of osteogenic markers in VSMCs cultured with control or osteogenic media. N= 4 independent experiments from 2 isolates. Significance was analyzed by Mixed-effects model test and q-values are shown. (E) The enrichment of ETS2 at the promoters of *BMP2*, *MSX2*, and *RUNX2* in VSMCs under osteogenic stimuli detected by Cut&Run-QPCR. N= 4 independent experiments from 2 isolates. Significance was analyzed by Mann-Whitney test and p-values are shown. (F) UCSC genome browser view shows BRD9 and BRG1 binding peaks at the promoter (red box), coding region (dosh line box), and enhancers (yellow boxes) of the *ETS2* locus in control VSMCs (black) or pre-osteogenic VSMCs (green). (G) The enrichment of BRD9, BRG1 and ETS2 at the *ETS2* promoter was detected by Cut&Run-QPCR. N= 6 (BRD9 and BRG1 independent experiments from 2 isolates. Significance was analysed by Mann-Whitney test and p-values are shown. (H) Detection of ETS2 protein expression in VSMCs cultured in control or osteogenic media with the presence or absence of BRD9 inhibitor. Cells were collected at individual stages of calcification progression and whole cell lysates were run on western blots to detect ETS2 and GAPDH. (I) KEGG analysis of ETS2-target genes (10588 genes) extracted from the published MotifMap database revealed associations with vascular smooth muscle contraction, regulation of actin cytoskeleton, and MAPK signalling pathway as top scoring terms, respectively. (J) MSigDB Hallmark analysis of ETS2 targets related to the function of vascular smooth muscle contraction showed associations with PI3K/AKT/mTOR pathway, inflammatory response, and KRAS signalling as top scoring terms, respectively. (K) 110 common genes between ncBAF-dependent genes and ETS2 targets in VSMCs. (L) Gene ontology analysis of 110 genes revealed associations with TNF alpha signalling via NFkB, Reactive Oxidative Species Pathway, and Hypoxia as top scoring terms, respectively.

### Inhibition of BRD9 modulates multiple cellular pathways activated during VSMC calcification

We next examined whether inhibiting BRD9 functionally regulates the above cellular pathways in VSMCs exposed to osteogenic stimuli. NFkB activation was detected by quantification of NFkB p52 activation in nuclear extracts. In pre-stage calcification, p52 activity was unchanged. In early-stage calcification the level of p52 activation was increased in pre-osteogenic cells and this activation was repressed by BRD9 depletion. In late-stage p52 was less active in calcified VSMCs than control cells and p52 activity was further downregulated by BRD9 depletion (Fig. S5A). This shows that NFkB activity increases as VSMC undergo phenotypic transition and BRD9 plays a role in promoting NFkB activation which may prime VSMCs to adopt an osteogenic phenotype. Using DAPI staining as an indicator of apoptosis we found the percentage of the sub-G1 population was increased in late-stage calcification compared to early-stage (Fig. S5B). I-BRD9 treatment reduced the sub-G1 population in both control and calcified cells in late-stage calcification (Fig. S5B), suggesting I-BRD9 treatment may inhibit apoptosis in VSMCs under osteogenic stress. We also examined glycolytic activity in control and calcified VSMCs in the presence or absence of I-BRD9. Glycolysis activity showed no dramatic changes in early-stage calcification (Fig S5C). In late-stage calcification, glycolysis activity, including glycolytic capacity and reserve capacity, were higher in control VSMCs than calcified VSMCs (Fig S5D). I-BRD9 treatment reduced glycolytic activity in both control and calcified VSMCs (Fig S5D), suggesting BRD9 may regulate VSMC metabolism. This evidence suggests that epigenetic influences of ncBAF complexes co-ordinately regulate diverse molecular pathways which promote VSMC calcification.

### The positive feedback loop by ncBAF-ETS2 in response to osteogenic stimuli in VSMCs

The transcription factor ETS2 was identified as a novel interactor of ncBAF complexes in pre-osteogenic VSMCs and as a target gene regulated by ncBAF in pre-osteogenic cells. This suggests the possibility of a positive feedback loop in pre-osteogenic VSMCs, where ncBAF upregulates ETS2, to form an activating complex that influences multiple cellular pathways upstream of VSMC calcification. To test this hypothesis, we validated the interaction between ncBAF and ETS2 in control or VSMCs exposed to osteogenic stimuli. IP-WB showed an enhanced interaction between ncBAF complexes and Ets2 in pre-osteogenic VSMCs compared to control (Fig 5A). Depletion of ETS2 (Fig S6A-B) showed that consistent with loss of BRD9 and reduction of BRG1, depleting ETS2 also attenuated VSMC calcification (Fig 5B-C). Silencing *ETS2* reduced the expression of *BMP2* and *RUNX2* in VSMCs in late-stage calcification (Fig 5D). Applying Cut&Run-QPCR, we confirmed that the occupancy of ETS2 at the promoters of *BMP2*, *MSX2* and *RUNX2* was increased in pre-osteogenic VSMCs compared to control (Fig 5E) suggesting ETS2 functions to promote VSMC calcification, in conjunction with BRG1 and BRD9.

To validate the function of ncBAF in regulating *ETS2* expression, we first showed the peak visualization of BRD9 occupancy at the promoter and coding region of *ETS2* was greater in pre-osteogenic VSMCs than control while BRG1 enrichment in the same regions was not as strong as BRD9 (Fig 5F). We also applied Cut&Run-QPCR to confirm that occupancy of BRD9 at the *ETS2* promoter was significantly increased in pre-osteogenic VSMCs with recruitment of BRG1 to the *ETS2* promoter again not as enriched as BRD9 (Fig. 5G). Interestingly, ETS2 and BRD9 were both enriched at the *ETS2* promoter in pre-osteogenic VSMCs (Fig. 5G), suggesting a positive feedback loop was established by BRD9-leading ncBAF complexes to stimulate ETS2 expression in response to mineral stress while ETS2 binding to ncBAF complexes acts to promote self-activation of ETS2. We next demonstrated that ETS2 protein was reduced in control and calcified VSMCs under treatment with I-BRD9 during calcification progression (Fig. 5H), indicating BRD9 is the major regulator of ETS2 activation.

We next explored genome-wide gene regulation by the ncBAF complex in cooperation with ETS2 during VSMC mineralization. Gene targets of ETS2 (10588 genes) were extracted from the published MotifMap database of transcription factor binding sites ^51,52^. KEGG pathway analysis of ETS2 targets showed vascular smooth muscle contraction as the most related cellular function (Fig 5I). To predict the possible biological role of ETS2 in VSMCs, we exported the gene list of ETS2 targets involved in vascular smooth muscle contraction and applied MSiDB Hallmark analysis. The result identified genes highly related to inflammatory response, KRAS signalling, and hypoxia (Fig. 5J), reiterating the list of ncBAF-dependent pathways in pre-osteogenic VSMCs (Fig. 4G).

110 of the 10588 ETS2 gene targets also showed differential peaks in the ncBAF CUT&RUN-Seq, indicating ETS2 and ncBAF complexes may coordinate at specific gene loci to remodel transcriptional patterns in VSMCs (Fig. 5K). Pathway enrichment analysis showed these 110 genes are related to the same diverse biological processes including positive regulation of NFkB signalling, reactive oxygen species pathway, and hypoxia (Fig. 5L), which were also identified by ncBAF-dependent genes in pre-osteogenic VSMCs (Fig. 4G). Taken together, these results demonstrate that in response to osteogenic stimuli, ncBAF upregulates ETS2, forming an activating complex that influences multiple cellular pathways that act to prime VSMCs for calcification.

### The co-activation of BRG1 and ETS2 define the pre-osteogenic phenotype of VSMCs in human artery single-cell RNA sequencing

To gain a better understanding of the relevance of this pathway in vascular calcification we examined ETS2 and ncBAF complexes *in vivo* in human artery samples using a published dataset of integrated single-cell RNA sequencing, which combined 22 scRNA-seq libraries to generate a comprehensive map of human atherosclerosis^10^. The smooth muscle cell (SMC) population was annotated by processing and transferring cell-type labels from the Tabula Sapiens (TS) vasculature single-cell atlas^53^, followed by re-clustering SMCs into 11 subclusters (Fig S7A). To characterize these SMC subclusters, we identified differentially expressed genes (DEGs) for each cluster and performed gene ontology (GO) analysis on cluster-specific DEGs to identify the overrepresented functions. Clusters 3, 5, 7, and 11 were classified as contractile SMCs (Fig. 6A) based on high expression of contractile markers (Fig. 6B) and enrichment of contractile GO terms (Fig. S7B), whereas clusters 0, 2, 4, and 9 exhibited GO enrichment for ECM remodelling with reduced expression of contractile markers and mild activation of osteogenic markers (Fig. 6B, Fig. S7B), indicating a transitional state toward an osteogenic phenotype and were therefore defined as transitional SMCs (Fig. 6A). Clusters 1 and 6 were defined as pre-osteogenic/ECM-remodelling SMCs (Fig. 6A) due to enrichment in ECM organization and ossification-related functions, accompanied by high expression of ECM markers, including collagens (Fig. 6B, Fig. S7B–D). In contrast, cluster 8 was classified as osteogenic/inflammatory SMCs (Fig. 6A) based on enrichment of ossification- and inflammation-related GO terms and high expression of osteogenic and inflammatory markers, including *RUNX2*, *SOX9*, *CCL2*, *CCL5*, *CCL19*, *CCL21*, and *CXCL12* (Fig. 6B, Fig. S7B, and E). An additional subcluster, cluster 10, was identified that showed enrichment of chemotaxis- and ECM-remodelling-related GO terms but lacked contractile or ossification-associated functions (Fig. S7B) and was therefore classified as “Other” SMCs. MSigDB Hallmark analysis of differentially expressed genes (DEGs) in clusters 1 and 6 revealed enrichment of apoptosis, glycolysis, and inflammatory response pathways (Fig. 6C–D). Notably, applying the same analysis to DEGs in cluster 8 showed enrichment of multiple inflammatory pathways, as well as apoptosis and glycolysis (Fig. 6E), closely aligning with the ncBAF-activated functions identified in pre-osteogenic VSMCs *in vitro*. We further confirmed that specific ncBAF target genes involved in the NF-κB activation pathway, including *CFLAR, PFKFB3, TMEM9B, and TNFAIP3*, identified by CUT&RUN sequencing (Fig. S4C), were highly expressed in osteogenic/inflammatory SMCs of cluster 8 (Fig. 6B), consistent with enhanced inflammatory activation. To determine whether these phenotypic changes in SMCs correlate with the progression of atherosclerosis, we performed pseudotime trajectory analysis across SMC clusters. The trajectory originated from contractile clusters 3, 5, 7, and 11, progressed through transitional clusters 0, 2, 4, and 9, and subsequently passed through clusters 1 and 6, which exhibit pre-osteogenic/ECM-remodelling phenotypes (Fig. 6F). The terminal state was cluster 8, characterized by osteogenic/inflammatory features (Fig. 6F), suggesting that ncBAF-regulated transcriptional programs drive phenotypic switching of VSMCs toward osteogenic and inflammatory states, thereby contributing to advanced stages of atherosclerosis.

**Figure 6.**
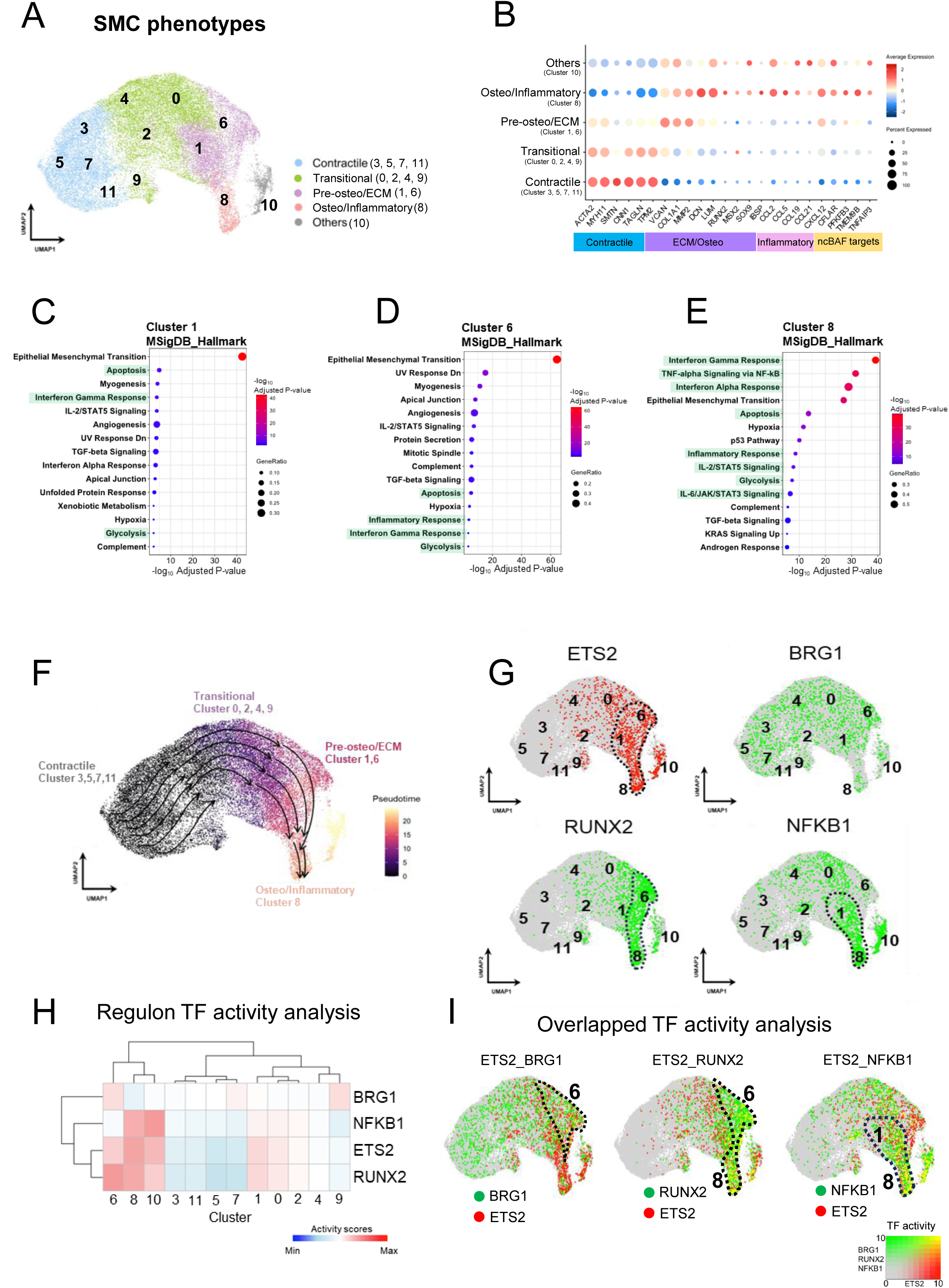
Active ETS2 coordinates with BRG1 to regulate the gene expressing patterns in VSMCs toward osteogenic/inflammatory phenotypes in the patients with atherosclerosis. (A) A UMAP plot shows the phenotypic annotations of SMCs by gene ontology analysis of DEGs. (B) A dotplot shows the gene expression levels of the specific markers as well as the gene targets of ncBAF complexes across all phenotypes of SMCs. (C) MSiDB Hallmark analysis of the DEGs in cluster 1 showed associations with apoptosis, interferon gamma response, and glycolysis which are included in the top 15 scoring terms. (D) MSiDB Hallmark analysis of the DEGs in cluster 6 showed associations with apoptosis, inflammatory response, interferon gamma response, and glycolysis which are included in the top 15 scoring terms. (E) MSiDB Hallmark analysis of the DEGs in cluster 8 showed associations with interferon gamma response, TNFα signalling via NFkB, interferon alpha response, apoptosis, and glycolysis which are included in the top 15 scoring terms. (F) UMAP embeddings shows the contractile-to osteogenic/inflammatory SMCs pseudotime trajectory calculated with Monocle3. SMC phenotypes for this analysis included contractile, intermediate, ECM-Producing/inflammatory, osteogenic/inflammatory SMCs. The trajectory root was defined as the contractile SMC clusters (cluster 3, 5, 7,11) in Fig 7A. (G) UMAPs present TF activity prediction of ETS2, BRG1, RUNX2, NFKB1 across all SMC clusters. (H) A heatmap presents the TF activity scores of BRG1, NFKB1, ETS2, and RUNX2 across all SMCs clusters. (I) UMAPs present the Transcription factor (TF) activity analysis showing SMCs with co-activation of ETS2 and BRG1 (ETS2_BRG1), ETS2 and RUNX2 (ETS2_RUNX2), and ETS2 and NFKB1 (ETS2_NFKB1).

Next, we mapped the transcription factor (TF) activity levels of ETS2 and BRG1, the ncBAF catalytic subunit, across all SMC clusters. SMCs with activated ETS2 were mainly enriched in clusters 1, 6, 8, and 10 (Fig 6G) which all showed activation of inflammation by GO term analysis. Clusters 1 and 6 appeared at an early stage and Cluster 8 at a late stage of pathological progression by pseudotime analysis (Fig 6F). In contrast, BRG1 was largely activated across different phenotypes of SMCs (Fig 6G), possibly due to BRG1 commonly existing in different subtypes of SWI/SNF complexes, which is consistent with distinct subtypes of SWI/SNF regulating phenotypic switching in VSMCs. The SMCs with ETS2-BRG1 co-activation were mainly located in cluster 6 (Fig 6H-I), indicating that ETS2-BRG1 coordination promotes pre-osteogenic phenotypes in SMCs in human vessels and this activation precedes the most pathological state observed in cluster 8 the final stage in the pseudotime trajectory.

Since ETS2-ncBAF complexes directly regulate osteogenic markers in pre-osteogenic VSMCs *in vitro*, we also investigated whether ETS2 and BRG1 are co-activated with the master osteogenic regulator RUNX2 *in vivo*. We first defined RUNX2 activity across all clusters and found it was highly activated in clusters 6 and 8 (Fig 6G), suggesting RUNX2 maintains activation from pre to final stages of pathological progression. Cluster 6 showed the highest level of RUNX2, ETS2, and BRG1 coordinated expression (Fig 6H-I), suggesting ETS2-BRG1 regulated transcriptional patterns correlate with expression of gene targets of RUNX2 and is an early event in pathological change. As the SMC clusters commonly co-displayed osteogenic and inflammatory phenotypes, we next examined NFKB1 activity and found it was highly activated in clusters 1, 8, and 10 (Fig 6G), which also overlap with ETS2-activated SMCs (Fig 6H-I). Taken together, this approach demonstrated that activating complexes composed of ETS2 and ncBAF drive early transcriptional changes that promote osteogenic/ECM and osteogenic/inflammatory phenotypes in VSMCs as they progress through the pathological stages of atherosclerosis to cluster 8 where both pathways intersect/overlap.

### Spatial distribution of ETS2 and ncBAF components in the human vasculature

We next investigated the spatial distribution of VSMCs exhibiting the phenotypes defined by our single-cell sequencing analysis and examined how this distribution correlated with the activation of Ets2–ncBAF complexes using spatial transcriptomic analysis of human carotid plaques from patients with atherosclerosis ^54^. Captured spots were first annotated based on transcriptional features of distinct cell types identified by single-cell RNA-sequencing analysis^10^, including endothelial cells (ECs), fibroblasts, SMCs, pericytes, macrophages, and plasmacytoid dendritic cells (pDCs) (Fig. 7A). We found that SMCs and macrophages were highly enriched within atherosclerotic plaques, with limited spatial overlap among these cell populations (Fig. 7A). Major SMC phenotypic states previously defined by single-cell RNA-sequencing analysis were also identified within the plaques based on their transcriptional similarities, including contractile, transitional, pre-osteogenic/ECM, and osteogenic/inflammatory SMCs (Fig. 7B). Notably, pre-osteogenic SMCs were predominantly located within SMC-enriched regions, whereas osteogenic SMCs primarily co-localized with macrophage-enriched areas of the plaques (Fig. 7C). These findings suggest that a macrophage-rich microenvironment may promote the differentiation of SMCs toward an overt osteogenic phenotype. Finally, we analysed the spatial expression patterns of *ETS2*, *BRG1*, and *BRD9* across all captured spots. Compared to *BRG1* and *BRD9*, the expression of *ETS2* was consistently found in both plaques (Fig. S8A).

**Figure 7.**
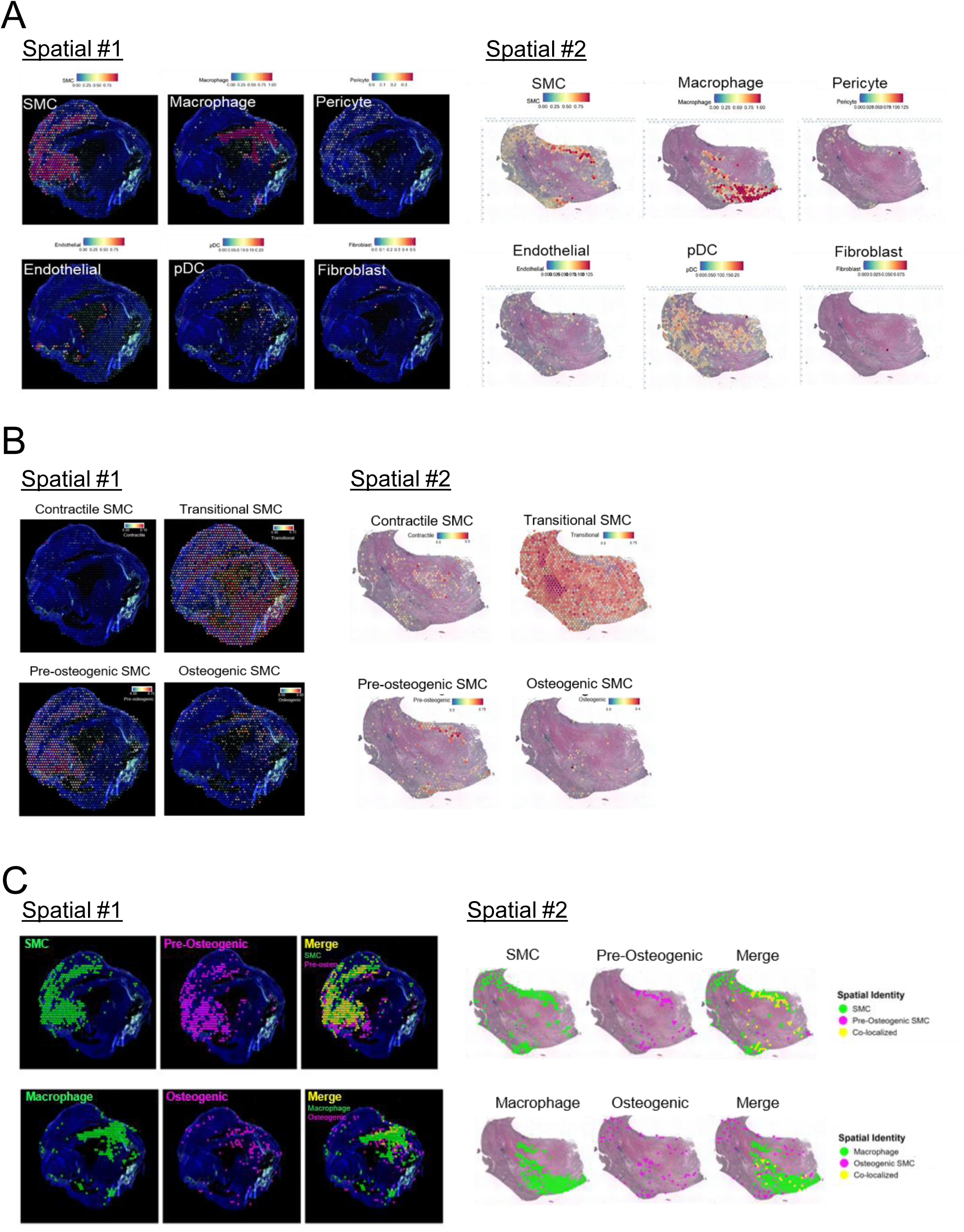
Spatial enrichment of osteogenic VSMC states in macrophage-rich regions of human carotid plaques and correlation of ETS2 with calcification severity. (A) Spatial transcriptomic feature plots from a carotid plaque sample Spatial #1 and Spatial #2 (10x Genomics Visium), showing the spatial distribution of major cell-type signatures including smooth muscle cell (SMC), macrophage, pericyte, endothelial cells, pDC, and fibroblasts across the tissue section. (B)Spatial maps of inferred VSMC transcriptional states in Spatial sample #1 and #2, including contractile, transitional, pre-osteogenic, and osteogenic programs, derived from single-cell signature scoring/deconvolution. (C) Top Panel: Spatial distribution of total Smooth Muscle Cells (SMCs) (green) and Pre-Osteogenic SMCs (magenta) across the tissue section. The merged images (right) highlight regions of Co-localization (yellow), indicating specific tissue niches where SMCs have initiated the transition toward a pre-osteogenic fate .Bottom Panel: Spatial overlap between Macrophages (green) and Osteogenic SMCs (magenta). The merged spatial plots demonstrate the proximity of macrophage infiltration to mature osteogenic SMC phenotypes, suggesting a potential inflammatory-driven mechanism for SMC phenotypic switching.

We next assessed ETS2 and BRG1 protein abundance *in vivo* by immunohistochemistry (IHC) in a cohort of human aortic samples. Von Kossa staining showed a positive correlation between age and vascular calcification (Fig. 8A-B). In parallel, the protein levels of ETS2 and BRG1 were positively correlated with each other (Fig. 8C), suggestive of coordinated upregulation of these factors in calcified vessels. To further define their relationship with calcification severity, we compared ETS2 and BRG1 staining intensities with the extent of Von Kossa staining. We found that ETS2, but not BRG1, was positively correlated with calcification levels (Fig. 8D-E), supporting ETS2 as a potential biomarker for calcification.

**Figure 8.**
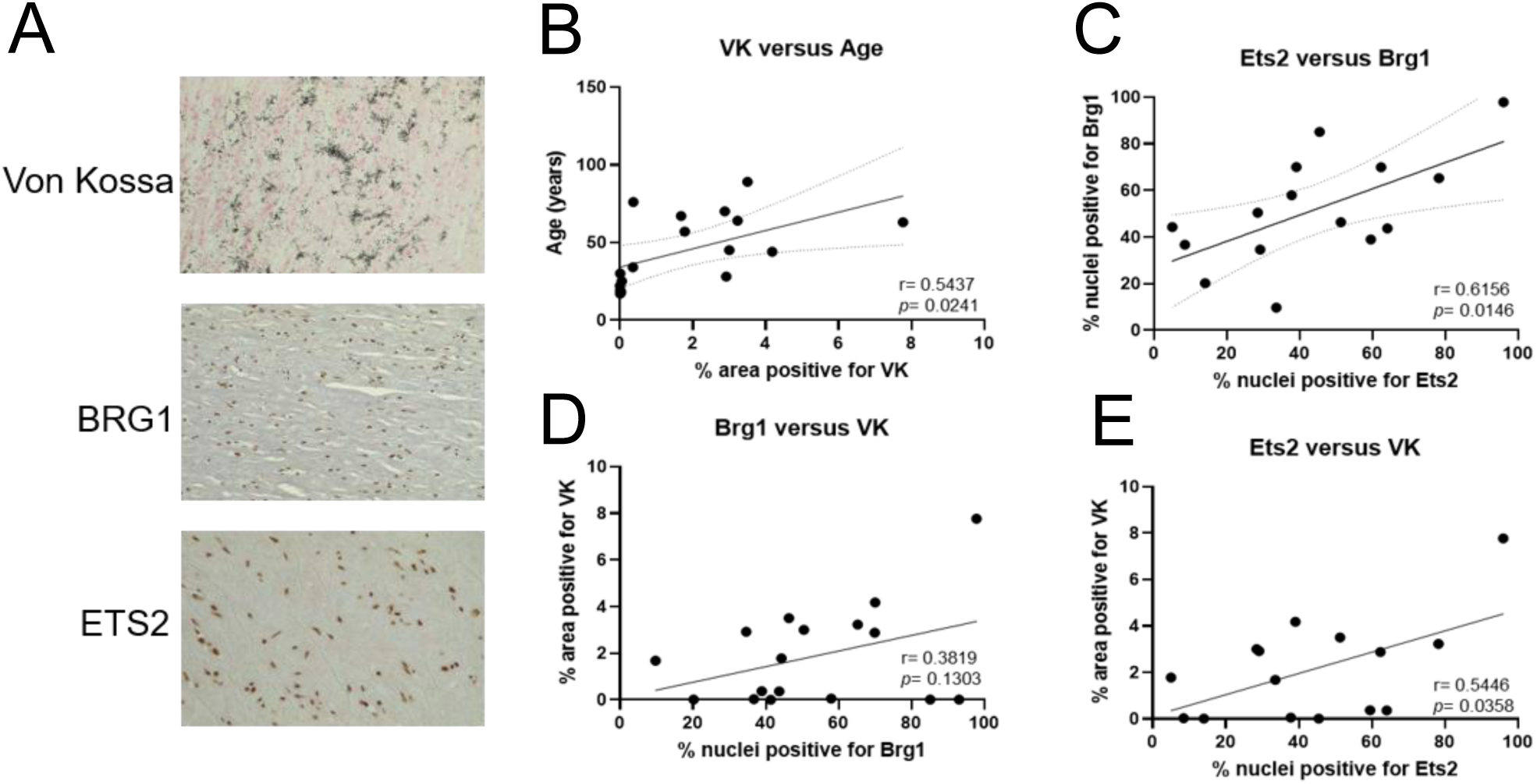
ETS2 protein levels correlated with calcification severity in human vessels supporting the potential clinical relevance of ETS2. (A) Representative staining of calcified vascular tissue showing mineral deposition by Von Kossa staining and immunohistochemical detection of BRG1 and ETS2 on adjacent sections. (B-E) Pearson correlation analyses (n = 16) showing relationships between calcification severity (Von Kossa–positive area) and age (B), ETS2-positive nuclei versus BRG1-positive nuclei (C), BRG1-positive nuclei versus calcification severity (D), and ETS2-positive nuclei versus calcification severity (E). Each dot represents one sample, and linear regression lines are shown. Pearson correlation coefficients (*r*) and *P* values are indicated in the plots.

**Figure 9.**
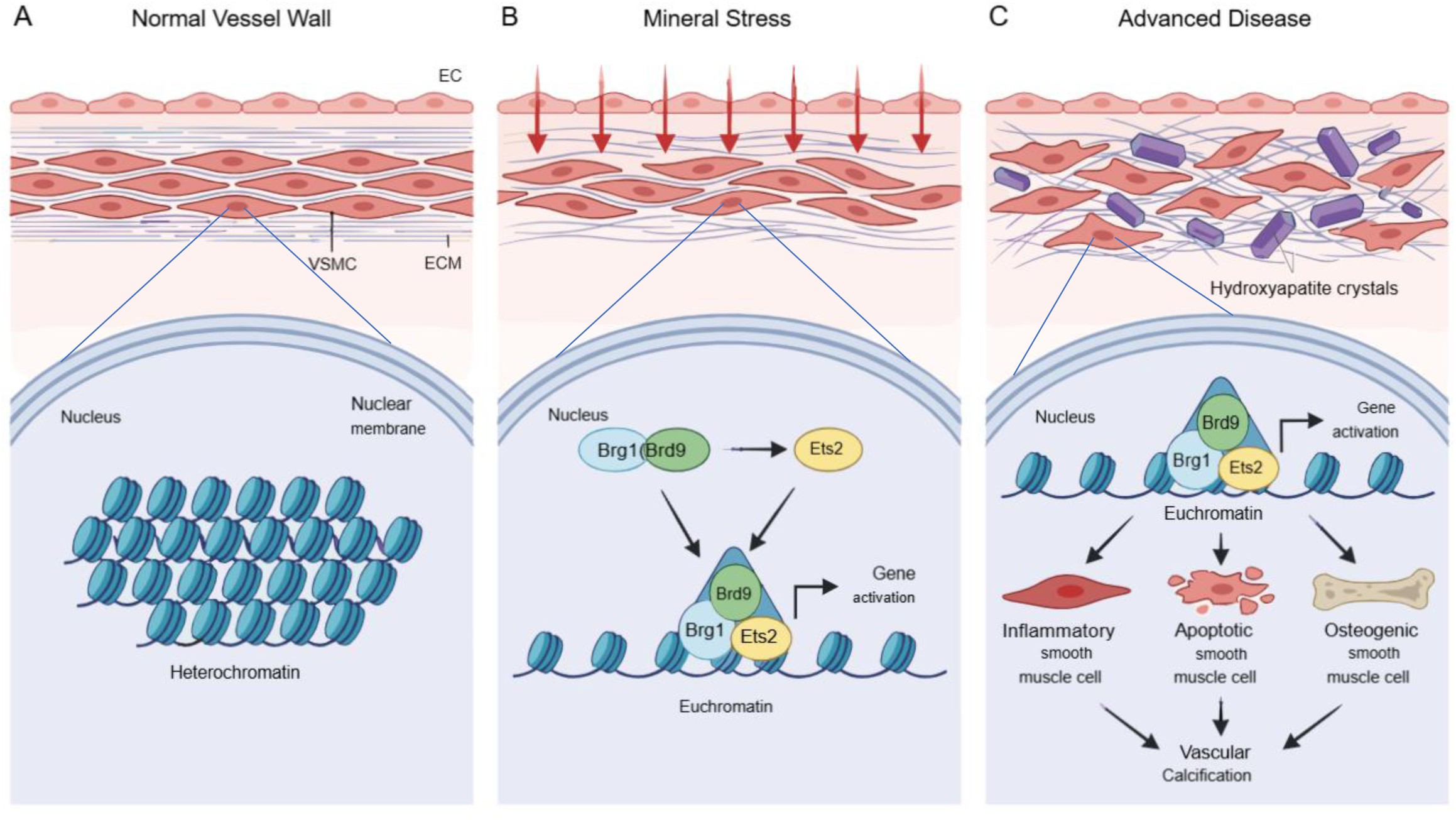
Graphic abstract: Mineral stress-induced epigenetic regulation of VSMC phenotypic switching in vascular calcification. A model of epigenetic remodelling and VSMC Fate During the progression of vascular calcification. (A) Normal Vessel Wall: Under physiological conditions, vascular smooth muscle cells (VSMCs) within the medial layer maintain a contractile phenotype. At the molecular level, chromatin is predominantly in a condensed heterochromatin state, restricting access to osteogenic and inflammatory gene loci. (B) Mineral Stress: chronic exposure to mineral stress (indicated by red arrows) triggers the activation of ncBAF chromatin remodelling complexes which regulates the transcription factor ETS2 expression. The assembly of BRG1, BRD9, and the transcription factor ETS2 leads gene activation on accessible euchromatin regions. This epigenetic “opening” allows for the initial activation of genes associated with phenotypic instability. (C) Advanced Disease: sustained gene activation through the ncBAF-ETS2 complex drives VSMCs transition into three pathological states: Inflammatory VSMCs: Contributing to a pro-inflammatory microenvironment. Apoptotic VSMCs: Providing niduses (vesicles) for further mineral deposition. Osteogenic VSMCs: Actively secreting bone-like matrix. Collectively, these cellular transformations culminate in progressive vascular calcification.

## Discussion

In this study, we identify the BRD9–BRG1 ncBAF complex as an early epigenetic regulator of the VSMC osteogenic transition and vascular calcification and show that it cooperates with the transcription factor ETS2 in pre-osteogenic VSMCs, to reshape transcriptional programs that coordinately promote multiple osteogenic and inflammatory responses. Notably, BRD9 also epigenetically upregulates ETS2 expression, establishing a positive feedback circuit that amplifies these programs during phenotypic switching. Consistent with these mechanistic findings, analysis of human artery single-cell datasets revealed co-activation of ETS2 and BRG1 in pre-osteo/ECM and osteo/inflammatory VSMC subsets with spatial transcriptomic profiling demonstrating regional clustering of these distinct SMC populations within atherosclerotic plaques. Immunohistochemistry showed that ETS2 levels correlated with calcification severity in human vessels supporting the potential clinical relevance of ETS2. Together, these results highlight stress-induced epigenetic priming as a critical early driver of VSMC plasticity and support targeting the ETS2–ncBAF axis as a potential therapeutic strategy for vascular calcification.

### Subunit switching within SWI/SNF complexes is required for cell plasticity of VSMCs

The ATPases BRG1 and BRM are two mutually exclusive catalytic subunits in SWI/SNF complexes. We showed that *BRG1* expression increased and *BRM* expression decreased during the progression of VSMC mineralization. We further confirmed that loss of BRG1 attenuated VSMC mineralization accompanied by reduced osteogenic marker expression. BRG1 occupied the promoters of osteogenic factors including *BMP2*, MSX2, and *RUNX2*, indicating the positive role of BRG1 in vascular calcification via its function in chromatin remodelling. Consistent with these findings, previous studies have revealed antagonistic roles for BRG1 and BRM in osteogenic differentiation in osteoblasts. Loss of BRG1 attenuated osteogenic differentiation but BRM-depletion showed accelerated progression to the mineralization phenotype^55^.

SWI/SNF complexes currently are categorised into three subtypes termed BAF, PBAF, and non-canonical BAF (ncBAF). We applied IP-MS to comprehensively identify compositional changes within SWI/SNF complexes specifically in VSMC calcification and demonstrated loss of BAF and increased PBAF and ncBAF complexes when VSMCs transform from contractile to osteogenic type. Importantly, we also determined that loss of BRD9 (a unique subunit of ncBAF) attenuated VSMC mineralization while depleting BRD7 and PBRM1 (unique subunits of PBAF) enhanced calcium deposition, indicating PBAF and ncBAF provide distinct epigenetic regulation in VSMCs in response to osteogenic stimuli. Previous studies in bone precursors have defined a role for other SWI/SNF complexes in mineralisation and are also suggestive of subunit switching. ARID2, a defined subunit of PBAF complexes, plays a similar role to BRG1 in promoting osteogenic differentiation^56^ while ARID1A and ARID1B which are mutually exclusive in BAF complexes differentially bind the osteocalcin (OSC) gene promoter in the pre- and late stages of osteogenic differentiation with ARID1A-BRM being replaced by ARID1B-BRG1^55^.

### Epigenetic influences by ncBAF complexes deregulated molecular pathways and led to a combinatorial effect in VSMC mineralization

We identified a panel of gene targets of ncBAF complexes in pre-osteogenic VSMCs.

Pathway enrichment analysis revealed that these genes are associated with pathways including TNFα signalling via NFκB activation, apoptosis, and glycolysis. Inhibition of BRD9 effectively modulated the activation of these pathways in VSMCs under osteogenic stress, suggesting that epigenetic influences by ncBAF complexes simultaneously activate multiple pathways that combined prime cells for phenotypic change in response to osteogenic stimuli. TNFα is a potent pro-inflammatory cytokine that is elevated in atherosclerosis, diabetes, and CKD. This cytokine creates an inflammatory environment that promotes osteogenic differentiation of VSMCs by upregulating various signalling pathways, including NFκB^57^. TNFα signalling involves the degradation of IκB proteins to release NFκB that then translocates to the nucleus to promote the transcription of target genes involved in inflammation and osteogenesis^58^. NFκB activation leads to the expression of pro-inflammatory genes such as IL-6, which further amplifies inflammatory signalling and promotes a pro-osteogenic environment^59^. Upregulated IL-6 expression has been shown in calcified VSMCs under mineral stress ^28^ and we demonstrated that inhibition of ncBAF complexes repressed elevated *IL-6* expression in late-stage calcified VSMCs (Fig 2C). A previous study showed BRD9 can activate interferon as well as NFkB signalling in response to TNFα during macrophage activation by bacterial endotoxin (lipid A)^60^. The same study showed inhibition of BRD9 repressed the expression of genes related to inflammatory pathways and this phenotype is consistent with our finding that inhibiting BRD9 repressed the genes relevant to NFkB activation in calcified VSMCs.

Damage and death of VSMCs play a significant role in vascular calcification. *In vitro* uraemia models of calcification are associated with significant increases in VSMC apoptosis^61^ with released apoptotic bodies forming a nidus for calcification^61–64^. Inhibition of apoptosis, by caspase inhibitors, significantly reduces both calcifying vesicle release and calcification^63^. Moreover, inhibition of apoptosis has been shown to attenuate calcification and conversely, stimulation of apoptosis increases the rate of calcification 10-fold^65^. In this study, we found that inhibiting BRD9 reduced the percentage of apoptotic cells in VSMCs treated with osteogenic media, indicating BRD9 regulates cellular apoptosis in VSMCs under calcification stimulation.

VSMCs exhibit high rates of glycolysis even under fully oxygenated conditions. PFKFB3, is a critical enzyme of glycolysis, that has been found to be upregulated in calcified VSMCs and arteries in the vitamin D_3_-induced calcification mouse model^50^. Silencing *PFKFB3* reduced the glycolysis rate and attenuated osteogenic differentiation in murine VSMCs^50^. In this study, genomic approaches pinpointed *PFKFB3* as an ncBAF-dependent gene in calcified VSMCs. Inhibiting BRD9 downregulated *PFKFB3* expression and reduced the glycolysis rate in both control and calcified VSMCs, suggesting ncBAF complexes contribute to maintenance of the glycolysis rate to support energy requirements in response to VSMC plasticity.

### Transcription factor ETS2 is defined as a novel regulator of VSMC calcification

ETS2 is a member of the DNA-binding domain ETS (E26 transformation-specific) family. A previous study showed that ETS factors such as ETS1 and ELK1 are involved in the activation of *RUNX2* gene transcription during osteoblast differentiation^66^. Our study for the first time demonstrated that ETS2 directly regulates osteogenic marker expression in VSMC calcification. We identified ETS2 as a novel interactor of ncBAF complexes in pre-osteogenic VSMCs, suggesting that ETS2 coordinates with ncBAF complexes, epigenetically regulating gene expression, to further mediate cellular pathways in response to osteogenic stimuli. Analysing the common gene targets between ncBAF and ETS2 showed they are highly related to the activation of NFkB signalling in pre-osteogenic VSMCs, indicating ETS2 contributes to phenotype switching towards an inflammatory phenotype in VSMCs. This observation is consistent with prior evidence showing that ETS2 functions to promote cytokine production in inflammatory macrophages^40^. Single-cell transcriptomic analysis of human atherosclerotic arteries further demonstrated that VSMCs with ETS2-BRG1 co-activation are associated with the early phenotypic transition towards a combined osteogenic and inflammatory state, while ETS2 activity persists in cells progressing toward high NFκB1 and RUNX2 activation. Pseudotime trajectory analysis showed that these VSMC clusters were enriched in more advanced stages of atherosclerosis, suggesting that ETS2 activation contributes to disease progression.

Spatial transcriptomics revealed that the most osteogenic VSMCs which also exhibited an inflammatory profile were enriched in macrophage-rich regions within human atherosclerotic plaques, suggesting a link between inflammatory microenvironments and osteogenic transition. In line with this, ETS2 staining intensity was positively correlated with calcification severity. Recent functional genomic studies identified ETS2 as the causal effector gene at the GWAS susceptibility locus chr21q22, where increased ETS2 expression drives inflammatory macrophage activation across multiple immune-mediated diseases ^40^. Together, these findings suggest that tight regulation of ETS2 is critical for human disease susceptibility, with both genetic variations affecting ETS2 regulation and aberrant ETS2 activation contributing to disease pathogenesis in a context-dependent manner.

We further demonstrated that pharmacological inhibition of BRD9 reduced ETS2 expression at both transcript and protein levels, indicating that BRD9 is required to maintain ETS2 homeostasis. These findings position BRD9 as a key upstream regulator of ETS2. The BRD9–ETS2 regulatory axis may represent a critical pathway linking chromatin remodelling to vascular pathology. Disruption of this axis could alter inflammatory signalling, metabolic programming, and VSMC plasticity—processes central to atherosclerosis and calcification progression. Collectively, these findings identify chromatin remodelling–dependent regulation of ETS2 as a key determinant of vascular cell state and suggest that targeting BRD9-dependent ETS2 regulation may represent a promising therapeutic strategy for cardiovascular disease.

## Acknowledgements

The proteomics was performed at the Denmark Hill Proteomics Facility. Sequencing was performed at the Genomics Research Platform, King’s College London. Graphical abstract and schematic were created with BioRender.com.

## Author Contributions

M.-Y. Wu, A. Durham and C.M. Shanahan contributed to the conception; M.-Y. Wu and C.M. Shanahan to experimental design; M.-Y. Wu, J. Thammaphet, S. Banday, S. Ahmad, C.-Y. Ho to the acquisition of data; M.-Y. Wu, J. Thammaphet, S. Banday, K Theofalatis, and C.M. Shanahan to analysis and interpretation of data. M.-Y. Wu and C.M. Shanahan wrote and revised the manuscript, and all authors provided final approval of the submitted version.

## SUPPLEMENTAL MATERIAL

### Expanded Materials and Methods

#### Cell Culture

Tissue samples were obtained with written informed consent for use in research, on a standard hospital consent form. Ethical approval for use of human cultures and tissue samples was approved by Local Research Ethics Committees all conforming to the principles outlined in the Declaration of Helsinki (NRES Committee London, REC No. LO/13/1950 and Great Ormond Street Hospital Institutional Review Board (12/LO/1186)). Human aortic vascular smooth muscle cell (VSMC) explant cultures were established from 3 individuals (both male and female), all healthy donors aged between 20-38 years (Supplementary Table 1).

Isolates were characterized for proliferative capacity and used at early passages (passages 8–13). VSMCs were maintained in M199 media supplied with 20% fetal bovine serum (FBS) and 1% penicillin/streptomycin/L-glutamate (PSG) in a humidified incubator with 5% CO_2_. To induce mineralization, VSMCs were treated with osteogenic media composed of M199 media with 5% FBS and 1% PSG, with additional calcium and phosphate buffer which is supplied by 180 mM CaCl_2_ and 100 mM NaH_2_PO_4_, to reach a final concentration of 2.7 mmol/L and 2.5 mmol/L, respectively. Calcium phosphate mineral deposition in VSMCs was visualized after cell fixation with 4% formaldehyde using Alizarin red (2%, wt/vol, pH 4.2) and quantified spectrophotometrically with cresolphthalein, as previously described^27,28^. Inhibitor I-BRD9 (SML1534, Sigma-Aldrich) was reconstituted in DMSO and VSMCs were treated with final concentrations of 5 µM in control or osteogenic media.

#### Small interfering RNA (siRNA)-mediated gene silencing

Commercially available human *BRD2*, *BRD3*, *BRD4*, *BRD7*, *BRD9*, *BRG1*, *PBRM1*, *ARID1A*, *ETS2* and non-targeting control siRNA (Dharmacon™) were used according to the manufacturer’s instructions. Briefly, 0.3 µl of 20 µM siRNA was mixed with 6 µl of HiPerFect Reagent (Qiagen) in 100 μL of serum-free media. siRNA mixtures were then incubated for 10 min at room temperature before dropping into cells. Cells were treated with 100 µl siRNA mixtures in the presence of 500 µl complete media supplemented with 5% FBS for 48 hours. Thereafter, cells were exposed to complete control media or osteogenic media to induce calcification. knockdown efficiency was confirmed by RT-QPCR and Western blotting.

#### Gene Expression Profiling and Quantitative Real-Time PCR

Total RNA was isolated from cells using RNA STAT-60 (CS-111, Amsbio) according to manufacturer’s protocol, followed by cDNA conversion using the M-MLV Reverse Transcriptase (M1701, Promega) according to the manufacturer’s protocol. Quantitative real-time PCR was performed in the ABI StepOne using SYBR™ Green PCR Master Mix (PB20.11-05, PCR Biosystems Ltd). The PCRs were performed in a 20μl reaction volume at 95°C for 10 minutes followed by 40 cycles at 95°C for 5 seconds and 60°C for 30 Seconds. Primers used are listed in Supplementary Table 2. Expression levels of genes of interest were normalized to 18S ribosomal RNA expression. Average fold differences were calculated by the comparative Ct method (2^-ΔΔCt^) between pairs of samples. For gene expression profiling of epigenetic regulators, RT2 Profiler PCR Arrays (Qiagen, PAHS-085ZA and PAHS-086ZA) was used according to manufacturer’s instructions.

#### General Procedure for Western Blotting

Cell pellets were lysed on ice for 30 minutes by lysis buffer (50 mM Tris HCl pH 7.5, 150 mM NaCl, 5mM EDTA pH 8, 0.5% Triton X-100 and protease inhibitors) and incubated on ice for 30 min, followed by 3 times sonication (5 seconds at power 1). Samples were spun down (20,000 g, 15 min, 4°C) and the supernatant was saved. Protein concentration was determined by DC™ Protein Assay Kit II (Bio-Rad, #5000112) and 30 μg whole cell lysates were loaded onto 4–15% precast polyacrylamide Gels (Bio-Rad #4561083) and separated by electrophoresis (1x running buffer containing 150 mM glycine, 20 mM Tris and 0.1 % SDS, 120 V, 60 min, at room temperature) and transferred to 0.2 μm PVDF membranes in transfer buffer (10% methanol, 150 mM glycine and 20 mM Tris) at 90 V, 120 min, 4°C. Membranes were blocked in Intercept (PBS) Blocking Buffer (LI-COR #927-70003) for 1 hour at room temperature, then washed three times with 1XPBST (1x PBS supplemented with 0.05% Tween20). Membranes were incubated with primary antibody overnight at 4°C, washed three time in 1XPBST, then incubated with secondary antibody (diluted 1:10000) for 1 hour at room temperature. Protein signals were detected by imaging on the Licor Odyssey CLx. The antibodies applied in this project are Brd9 (1:1000, ab259839, Abcam), Brg1(1:5000, ab110641, Abcam), Pbrm1 (1:5000, A301-591A, Bethyl Laboratories), Ets2 (1:1000, PA5-28053, Invitrogen), Gapdh (1:5000, ab9484, Abcam), β-actin (1:5000, A228, Sigma).

#### Immunoprecipitation Followed by Mass Spectrometry analysis

VSMC pellets fresh or snap frozen were used for immunoprecipitation. 2 μg antibody (Brd9-Abcam ab259839, Brg1-Abcam ab110641, Rabbit IgG-Cell Signaling Technology #2729) was cross-linked to Dynabeads Protein A (Invitrogen, # 10001D) using dimethyl pimelimidate (DMP, Sigma-Aldrich, #D8388). Cells were lysed on ice in lysis buffer (50 mM Tris HCl pH 7.5, 150 mM NaCl, 5mM EDTA pH 8, 0.5% Triton X-100 and protease inhibitors). 750 μg of protein lysate was incubated with crosslinked beads for 16 hrs at 4 °C. Immunoprecipitations were washed 3 times with wash buffer (50 mM Tris HCl pH 7.5, 150 mM NaCl) and eluted in 50 ul of 7.5% SDS, 20 mM DTT in lysis buffer boiled at 95 degrees for 5 minutes. Take 20 ul of elutes to run Western blotting detection. 30 ul of elutes then was alkylated for 1 hour with 40 mM iodoacetamide at 25°C in the dark and proteins were cleaned up using the SP3 beads (VWR, Sera-Mag Speed Beads 1:1, CAT# 09-981-121, 09-981-123, rinsed with water). SP3 beads mix (20 μg/μl) was added to the alkylated IP sample (1 μg/μl). Sample was acidified to pH 2-3 using 10% formic acid and bound to SP3 beads with one volume 100% acetonitrile. Beads were rinsed twice with 70% ethanol followed once by 100% acetonitrile. Proteins were disassociated from SP3 beads by 50 ul of 50 mM triethylammonium bicarbonate (TEAB) buffer pH 8.5 and digested with 1 μg trypsin (Pierce) per sample overnight at 37 °C. SP3 beads were not removed for peptide clean up. Resulting peptides were bound to the SP3 beads by addition of one volume acetonitrile and washed twice with 100% acetonitrile. After drying, clean peptides were eluted by using 2% DMSO in water. Clean peptides were vacuum dried and sent to mass spectrometry analysis. Samples were resuspended in 2% (v/v) acetonitrile, 0.05% (v/v) formic acid and peptides were resolved by reversed phase chromatography on a 75µm C18 Pepmap column (50cm length) using a three-step linear gradient of 80% acetonitrile in 0.1% formic acid (U3000 UHPLC NanoLC system; Thermo Fisher Scientific, UK). The gradient was delivered to elute the peptides at a flow rate of 250nl/min over 60 min starting at 5% B (0-5 minutes) and increasing solvent to 40% B (5-40 minutes) prior to a wash step at 99% B (40-45 minutes) followed by an equilibration step at 5% B (45-60 minutes). The eluate was ionised by electrospray ionisation using an Orbitrap Fusion Lumos (Thermo Fisher Scientific, UK) operating under Xcalibur v4.4.16.14. The instrument was first programmed to acquire using an Orbitrap-Ion Trap method by defining a 3s cycle time between a full MS scan and MS/MS fragmentation. Orbitrap spectra (FTMS1) were collected at a resolution of 120,000 over a scan range of m/z 375-1500 with an automatic gain control (AGC) setting of 4.0e5 with a maximum injection time of 35 ms. Monoisotopic precursor ions were filtered using charge state (+2 to +7) with an intensity threshold set between 5.0e3 to 1.0e20 and a dynamic exclusion window of 35 secs ± 10 ppm. MS2 precursor ions were isolated in the quadrupole set to a mass width filter of 1.6 m/z. Ion trap fragmentation spectra (ITMS2) were collected with an AGC target setting of 1.0e4 with a maximum injection time of 35 ms with CID collision energy set at 35%.

Data were processed using Proteome Discoverer (v2.5; Thermo Fisher Scientific) to search against Uniprot Swiss-prot Human Taxonomy (50,442 entries) with Mascot search algorithm (v2.6.0; https://www.matrixscience.com) and the Sequest search algorithm^30^. Precursor mass tolerance was set to 20 ppm with fragment mass tolerance set to 0.6 Da with a maximum of two missed cleavages. Variable modifications included: Carbamido-methylation (Cys) and Oxidation (Met). Database generated files (.msf) were uploaded into Scaffold software (v 5.0.0; https://www.proteomesoftware.com) for visualisation of fragmentation spectra. The differences between means for protein abundance of control and pre-osteogenic VSMCs were calculated using label-free quantifying (LFQ) intensities and plotted versus log_10_ *P value*s determined by two-tailed t test (Perseus Software, v.1.6.1.3). Protein-Protein interaction network analysis was processed by using STRING database 2023^34^.

#### Quantitative NFκB activation

Quantitative NFκB activation was performed by NFκB p52 Transcription Factor Assay Kit (Colorimetric) (ab207219, Abcam) according to the manufacturer’s protocol. Briefly, 5 ug nuclear extract from VSMCs transfected with non-targeting or targeting siRNA culturing in control or osteogenic media, was diluted in 30 ul Complete Binding Buffer (CBB buffer) provided by the kit then loaded into 96-well NFκB assay plate which is pre-coated with an oligonucleotide containing NFκB p52 consensus binding site followed by incubation for 1 hour at room temperature with mild agitation. Washing wells 3 times with 200 ul 1x wash buffer. After that, adding 100 ul p52 antibodies (1:1000 in 1x Antibody Binding Buffer) and incubate for 1 hour at room temperature. Add 100 ul secondary antibodies (1:1000 in 1xAntibody Binding Buffer) and incubate for 1 hour at room temperature. After washing wells 4 times, add 100 ul developing solution and incubate up to 5 minutes at room temperature, followed by adding 100 ul stop solution. Absorbance was read on a spectrophotometer at OD 450 nm within 5 minutes after adding stop solution with a reference wavelength of OD 655 nm. Blank the plate reader according to the manufacturer’s instructions using the blank wells. Absorbance of each datapoint was performed by normalization to the control condition.

#### Quantification of the percentage of apoptotic cells

1*10^6^ cells in 0.5 ml 1xPBS were fixed by mixing with 4.5 ml of ice-cold 70% ethanol. Keep the cells in the fixative for at least 12 hours at 4 degrees. After that, centrifuge ethanol-suspend cells for 5 minutes at 500 *g* and decant ethanol thoroughly. Wash cells once with ice-cold 1xPBS then resuspend cell pellet in 0.5 ml of DAPI solution (1 ug/ml DAPI, 0.1%Triton X-100 in 1xPBS), incubating for 5 minutes at 37 degrees. Cellular fluorescence is quantified by NucleoCounter® NC-3000™ instrument and apoptotic cells with fragmented DNA are seen and quantified as a sub-G1 peak in a DNA content histogram.

#### Glycolysis stress test by Seahorse analysis

VSMCs were seeded into Seahorse XFe24 cell culture microplates at a density of 4*10^4^ cells/well. VSMCs were cultured in control or osteogenic media in the presence or absence of BRD9 inhibitor. Glycolysis stress test was performed when cells reached early- or late-stage calcification. Hydrate a sensor cartridge in Seahorse XF Calibrant at 37 °C in a non-CO_2_ incubator overnight before Glycolysis stress test. On the day of analysis, cells were washed 3 times with Seahorse XF assay medium and incubated in a CO_2_-free incubator at 37°C for 45 minutes. The ECAR (acidification rate) assay was performed on the Seahorse XF Analyzer, and the ECAR values were normalized using cell intensity acquired by DRAQ5 staining. The relative levels of glycolysis and glycolytic capacity were calculated based on ECAR data obtained in the glycolysis stress tests. Inhibitors and activators were used at the following concentrations: glucose (10 mM), oligomycin (1 uM), and 2-DG (50 mM).

#### Cut&Run Sequencing and analysis

Cut&Run chromatin IP was performed using a centrifugation-based protocol as previous published^41^. Two million cells were harvested by centrifugation (600 g, 3 min) and washed in ice cold phosphate-buffered saline (PBS). Nuclei were isolated by hypotonic lysis in 1 ml NE1 buffer (20 mM HEPES-KOH pH 7.9, 10 mM KCl, 0.5 mM spermidine, 0.1% Triton X-100, 20% Glycerol) for 5 min on ice followed by centrifugation. Nuclei were briefly washed in 1.5 ml Buffer 1 (20 mM HEPES pH 7.5; 150 mM NaCl; 2 mM EDTA; 0.5 mM Spermidine; 0.1% BSA) and then washed in 1.5 ml Buffer 2 (20 mM HEPES pH 7.5; 150 mM NaCl; 0.5 mM Spermidine; 0.1% BSA). Nuclei were resuspended in 500 µl Buffer 2 and 2 ug of antibodies (Brd9-Abcam ab259839, Brg1-Abcam ab110641, H3K27ac-Abcam ab4729, Rabbit IgG-Cell Signaling Technology #2729) were added and incubated at 4°C for 2.5 hrs. Nuclei were washed 3 times in 1 ml Buffer 2 to remove unbound antibody. Nuclei were resupended in 300 µl Buffer 2 and 5 µl pA-MNase added and incubated at 4°C for 1 hr. After washing out unbound pA-MNase, nuclei were resuspended in Buffer 2 and quickly added 100 mM CaCl_2_ to reach a final concentration of 2 mM to activate MNase for 15 mins at 0 degree. The reaction was quenched by adding a master mix of EDTA (10 mM) and EGTA (20mM) on ice for 10 minutes. Cleaved fragments were liberated into the supernatant by incubating the nuclei at 4°C for 30 minutes, and nuclei were pelleted by centrifugation as above. DNA fragments were purified by proteinase K digestion followed by Phenol-chloroform-isoamyl alcohol mixture (PCI-mixture) extraction and used for RT-QPCR or the construction of sequencing libraries. Library preparation was performed using the NEBNext Ultra II DNA Library Preparation Kit (NEB#7645S) and sequenced using Illumina NextSeq2000 with p2 flow cell 100 cycle kit providing an estimated reads of 33 million per sample. Size distribution of the libraries was checked using high sensitivity D1000 ScreenTape in Agilent 4150 TapeStation system and quantitation was performed in Qubit Fluorometer (Invitrogen).

Bowtie2 package was applied for reads alignment against reference genome human hg38 assembly and saved in bam format. Bam files then were converted into bigwig files using bamCoverage with normalization method as normalizing to reads per kilobase per million (RPKM). Bigwig files were applied for peak visualization using UCSC Genome Browser https://genome.ucsc.edu/. Bam files were converted to Bed files using Bedtools package followed by peak calling using MACS2 software using default parameters with a cutoff of q-value equal to 0.05 and saved in narrowPeak format. Differential peaks between different conditions (control and pre-osteogenic VSMCs) were identified by DiffBind package using narrowPeak bed files and Bam reads files from 2 biological repeats of each condition. The output Bed file of DiffBind analysis was applied for ChIPseeker software with gencode.v38. annotation.gtf file for peak annotation and visualization. Heatmaps for score distribution across genomic regions were plotted by matrix files being generated by ComputeMatrix package using region narrowPeak bed files and score bigwig files. All data has been deposited in Gene Expression Omnibus (GEO) under accession number (GSEXXXXXX).

#### RNA Sequencing and analysis

Total RNA was isolated from control or pre-osteogenic VSMCs using Direct-zol^TM^ RNA Miniprep kit (Zymo Research) according to the manufacturer’s instructions. Library was created with 1 ug of total RNA and the preparation was performed using Illumina Stranded mRNA Prep, Ligation kit with IDT-ILMN DNA/RNA UD Index Set A and sequenced using Illumina NextSeq2000 with p2 flow cell 100 cycle kit. Quality of RNA-seq libraries were checked using high sensitivity D1000 ScreenTape in Agilent 4150 TapeStation system and quantitation was performed in Qubit Fluorometer (Invitrogen). HISAT2 package was applied for reads alignment against reference genome human hg38 assembly and generated Bam files for quantifying transcripts using StringTie software with GENCODE annotation (gencode.v38.annotation.gtf). The output files generated by StringTie package were applied for DEseq2 package to identify the differential gene expressions across the conditions of interest. Differentially expressed genes were identified using the DEseq2 package and p value adjusted for multiple testing with the Benjamini-Hochberg procedure which controls false discovery rate (FDR) and genes with adjusted p-values <0.05 were considered significant. Pathway enrichment analysis of genes that fulfilled these criteria was performed using the Enrichr software (http://amp.pharm.mssm.edu/Enrichr/)^42,43^. FeatureCounts package is applied for counting mapped reads for genomic features such as genes, exons, promoters, or gene bodies.

#### Pathway enrichment and String interaction network analysis

STRING interaction network analysis^34^ was applied to define potential gene regulation and protein-protein interaction. Pathway enrichment analysis with the Human Molecular Signatures Database (MSigDB) and Gene Ontology analysis are applied to identify specific changes in pathway activation in the conditions of interests using the Enrichr software (http://amp.pharm.mssm.edu/Enrichr/)^42,43^.

#### Analysis of public human atherosclerotic scRNA-seq data

Integrated human single-cell RNA sequencing (scRNA-seq) data derived from atherosclerotic arteries, as reported previously¹⁰, were utilized in this study. Cells annotated as “SMC” in the level 1 classification of the integrated dataset were subsetted for downstream analysis. The subsetted SMC population was re-clustered using the first 30 principal components (PCs) with a clustering resolution of 0.6. Differentially expressed genes (DEGs) for each SMC cluster were identified using the FindAllMarkers function in Seurat, applying the Wilcoxon rank-sum test. Genes were filtered using logfc.threshold = 0.25 and min.pct = 0.1. SMC phenotypes were inferred based on canonical marker gene expression combined with Gene Ontology (GO) enrichment analysis. GO Biological Process (GO:BP) enrichment analysis was performed using the clusterProfiler package (v4.14.0). The top 500 DEGs per cluster, ranked by log2 fold change, were used as input. Significantly enriched biological processes (adjusted P-value < 0.05) were used to support phenotypic annotation of SMC clusters.

#### Pseudotime cell trajectory analysis for SMCs

Pseudotime analysis of single-cell transcriptomic data from human datasets was performed using Monocle3 (v1.3.7). Seurat objects were converted to Monocle3 CellDataSet objects using the as.cell_data_set function, with cells grouped according to Seurat clusters or annotated SMC phenotypes. Dimensionality reduction was performed using UMAP, followed by clustering with the Leiden algorithm using a resolution parameter of 1 × 10⁻⁴. Trajectory inference was conducted using the learn_graph function. Gene annotation information was incorporated into the CellDataSet object prior to downstream analyses. The contractile SMC phenotype was designated as the root state, and pseudotime ordering was performed using the order_cells function.

#### Transcription factor (TF) activity inference

Transcription factor activity was inferred from human scRNA-seq data using the decoupleR package (v2.12.0). TF–target interaction networks were obtained from the CollecTRI resource, a curated collection of transcriptional regulatory interactions. TF activity scores were estimated using the Univariate Linear Model (ULM) method, which evaluates the enrichment of TF target gene weights at the single-cell level. Mean TF activity scores were calculated for each cluster or SMC phenotype and ranked in descending order. TFs with statistically significant activity (P-value < 0.05) were retained for downstream analysis and interpretation.

#### Spatial transcriptomics Data preprocessing

Spatial transcriptomics data from human carotid atherosclerotic plaques, as reported previously⁵⁴, were retrieved from Gene Expression Omnibus (GSE241346). These data were re-analyzed using Seurat (v4.4.0) in R (v4.2.3). Gene expression count matrices were obtained from 10x Genomics Visium output files and used to generate individual Seurat objects. Standard quality control criteria were applied, including a minimum threshold of 200 detected features per spot. High-resolution histological images and spatial barcode coordinates were imported and aligned with the RNA assay. Only barcodes corresponding to retained expression data were included. Spatial images were incorporated as slice objects for downstream visualization. The two samples were merged into a combined dataset. To reduce technical variability and improve clustering performance, normalization was performed using SCTransform independently for each sample. Quality control metrics, including total RNA counts (nCount_RNA), were assessed using spatial distribution plots and violin plots. Spots with zero RNA counts were excluded from further analysis.

#### Dimensionality reduction and clustering of spatial data

Dimensionality reduction and clustering were performed using the standard Seurat workflow. Principal component analysis (PCA) was conducted using the top 30 PCs. A shared nearest neighbor graph was constructed, followed by clustering using the Leiden algorithm with a resolution of 0.3. UMAP was used for visualization of low-dimensional embeddings. Cluster identities were visualized using both UMAP and spatial feature plots. Expression patterns of selected SMC marker genes and genes of interest were examined to assess spatial localization and regional enrichment.

#### Reference-based cell type mapping

To map single-cell phenotypes onto spatial transcriptomics data, a reference scRNA-seq dataset derived from human vascular tissue¹⁰ was re-analyzed and curated to define SMC subtypes. Cell-type annotations were refined based on canonical marker genes and biological relevance, focusing on SMC phenotypes including contractile, transitional, pre-osteogenic, and osteo/inflammatory states. The reference dataset was normalized using SCTransform, followed by PCA and UMAP embedding. Anchor-based integration between the reference scRNA-seq dataset and spatial transcriptomics data was performed to identify shared transcriptional profiles. Cell-type probabilities were transferred using a PCA-weighted label transfer approach and stored in a dedicated assay for downstream analysis.

#### Spatial annotation and visualization

Spatial distribution of predicted SMC phenotypes was visualized by plotting deconvolved phenotype probabilities across the tissue sections. Feature plots of inferred cell states were generated using customized visualization parameters to enhance clarity and contrast. All spatial plots were exported as high-resolution images at 600 dpi for publication-quality rendering. All computational analyses were performed in R (version 4.2.3) using Seurat (version 4.4.0), sctransform (version 0.3.5), patchwork (version 1.1.3), and ggplot2 (version 3.4.4).

#### Immunohistochemistry

Formalin fixed paraffin embedded (FFPE) sections of human arteries (7μm thickness) were cleared in xylene and rehydrated through graded ethanol. Antigen retrieval was performed via sodium citrate (pH 6, #SK4100) or Tris-EDTA (pH 9) according to antibody specification. Sections were treated for 20 min with 3% hydrogen peroxide in methanol to quench endogenous peroxidase. Next, sections were blocked in 10% horse or goat serum and primary antibodies against Ets2 (1:200, OriGene Catalog # TA505372), Brg1 (1: 200, Abcam ab110641) were diluted in blocking buffer and incubation was performed overnight at 4°C. Control sections were incubated without primary antibody. Detection was via the avidin-biotinylated-HRP complex (ABC) method using an Elite ABC kit (Vector Laboratories). Visualization of bound HRP was performed using a reaction with 3-3’-diaminobenzidine (DAB) (Vector labs #SK-4100) and the reaction was stopped by incubating the sections in distilled water. Nuclei were counterstained with hematoxylin. Slides were then dehydrated in graded ethanol, cleared in xylene and mounted using DPX mounting medium (Sigma). Images were captured using a Leica brightfield microscope and processed using GIMP2. Nuclei were counted using Image J software. Operators were blinded to the identity of all samples analysed. Picrosirius red and Alcian blue staining were performed to evaluate status of the extracellular matrix and development of fibrosis. Von Kossa staining was used to detect calcium deposition in the vessel wall25. Briefly, 7μm thick FFPE sections were deparaffinized in xylene and rehydrated in different grades of alcohol. The sections were then incubated with 1% Silver Nitrate solution under a 100-watt light bulb for 1.5hrs. The unreacted silver nitrate was removed 5% sodium thiosulfate solution for 5 min following several washes with distilled water. A nuclear fast red solution was used to stain nuclei and cytoplasm. The slides were then dehydrated through graded alcohols and cleared in xylene before coverslips were attached using DPX mounting media (Sigma). The collagen network was highlighted by picrosirius red staining. Sections were incubated in 0.1% picrosirius red solution for 15 min and quickly differentiated in 2 baths of acidified water before dehydration and mounting. Alcian blue staining was performed to visualize the accumulation of glycosaminoglycans (GAGs) in cells. Sections were incubated in alcian blue solution for 30 min then rinsed in distilled water before dehydration and mounting according to manufacturer’s instructions (Merck Life Sciences).

#### Statistical Analysis

Quantitative data are presented as mean±SEM or as the median and interquartile range as appropriate. All data were analysed with GraphPad (Prism 10.1) statistical software. Normality of data was tested by the Shapiro-Wilk test and the Kolmogorov– Smirnov test. For comparison of two independent groups, the parametric unpaired t-test was used. The Mann-Whitney test was used where the distribution of data points failed the normality test. For comparison of multiple groups, ordinary one-way ANOVA or Mixed-effects model test was used and the p-value shown for significant difference. Where the numbers of data points were less than ten the Krukal-Wallis test or mixed model analysis was performed and q values were adjusted for multiple testing with the two-stage step-up method of Benjamini, Krieger and Yekutieli false discovery rate (FDR) correction. The p-values and q-values are presented with numeric values using scientific format when less than 0.0001.

---

**Supplementary Figure 1:**
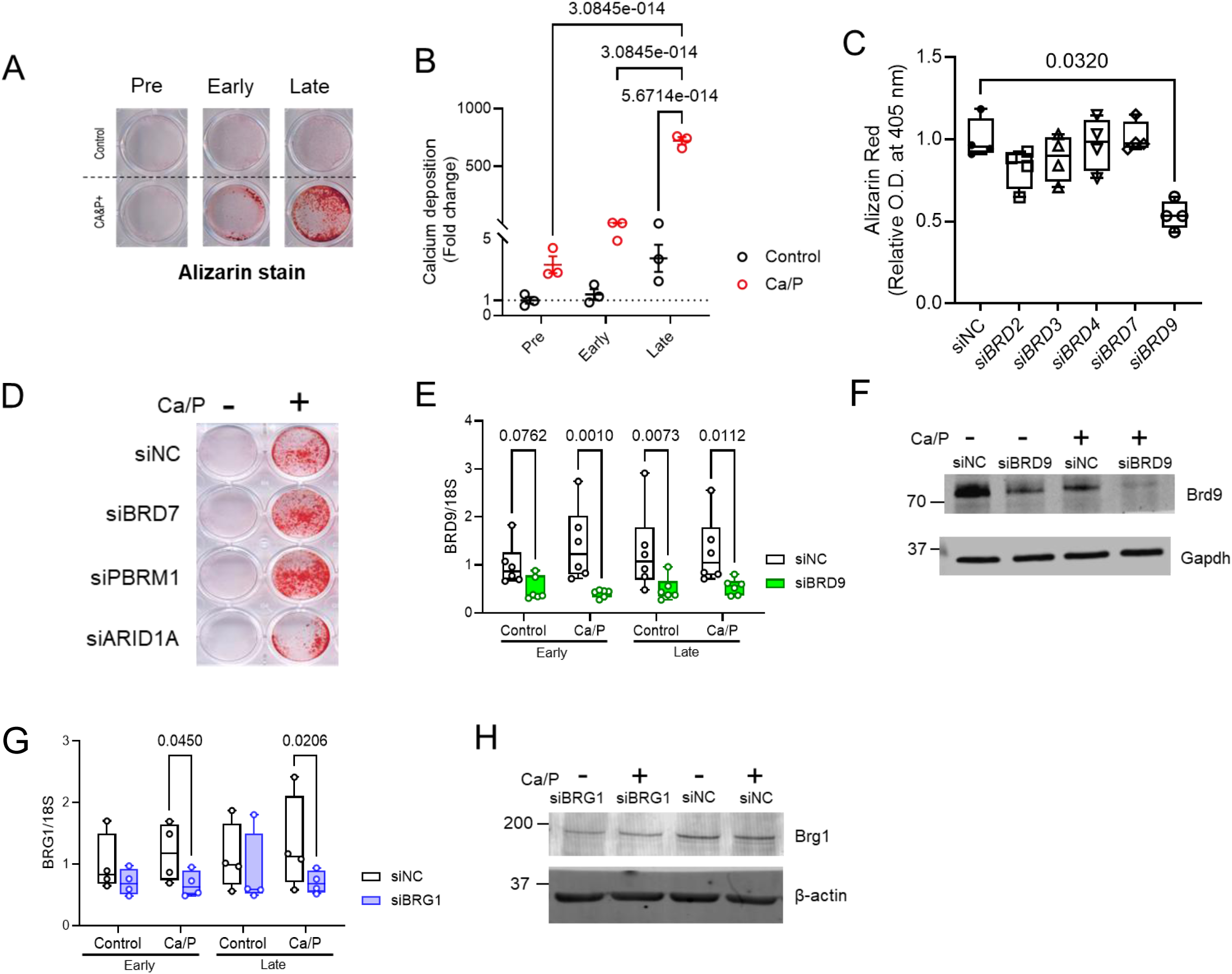
(A) Alizarin red staining of control and calcified VSMCs at the indicated calcification stages. (B) o-Cresolphatalein assay was used to quantify calcium content of control or calcified VSMCs for the indicated calcification stages. Total calcium content was normalised to total protein content in each sample. N= 3 independent experiments from 3 isolates. (C) Quantified Alizarin red staining of VSMCs after depletion of different Bromodomain protein family members and treatment with control (Control) or osteogenic media to induce calcification. (D) Alizarin red staining of VSMCs cultured in control or osteogenic media to induce calcification after siRNA depletion of Brd7, Pbrm1, Arid1a, and control non-targeting siRNA (siNC). (E) Confirmation of *BRD9* knockdown efficiency by QPCR. VSMCs were transfected with non-targeting siRNA (siNC) and siRNA targeting *BRD9* (siBRD9) and cultured in the presence or absence of osteogenic stimuli (Ca/P). Samples were collected at the early- and late-stage of calcification and the expression of *BRD9* was detected by QPCR. N= 6 independent experiments from 3 isolates. Significance was analyzed by Mixed-effects model test and q-values are shown. (F) Confirmation of *BRD9* knockdown efficiency by Western blotting. (G-H) Confirmation of *BRG1* knockdown efficiency in the presence or absence of osteogenic stimuli (Ca/P) by QPCR (G) and Western blotting (H).

**Supplementary Figure 2:**
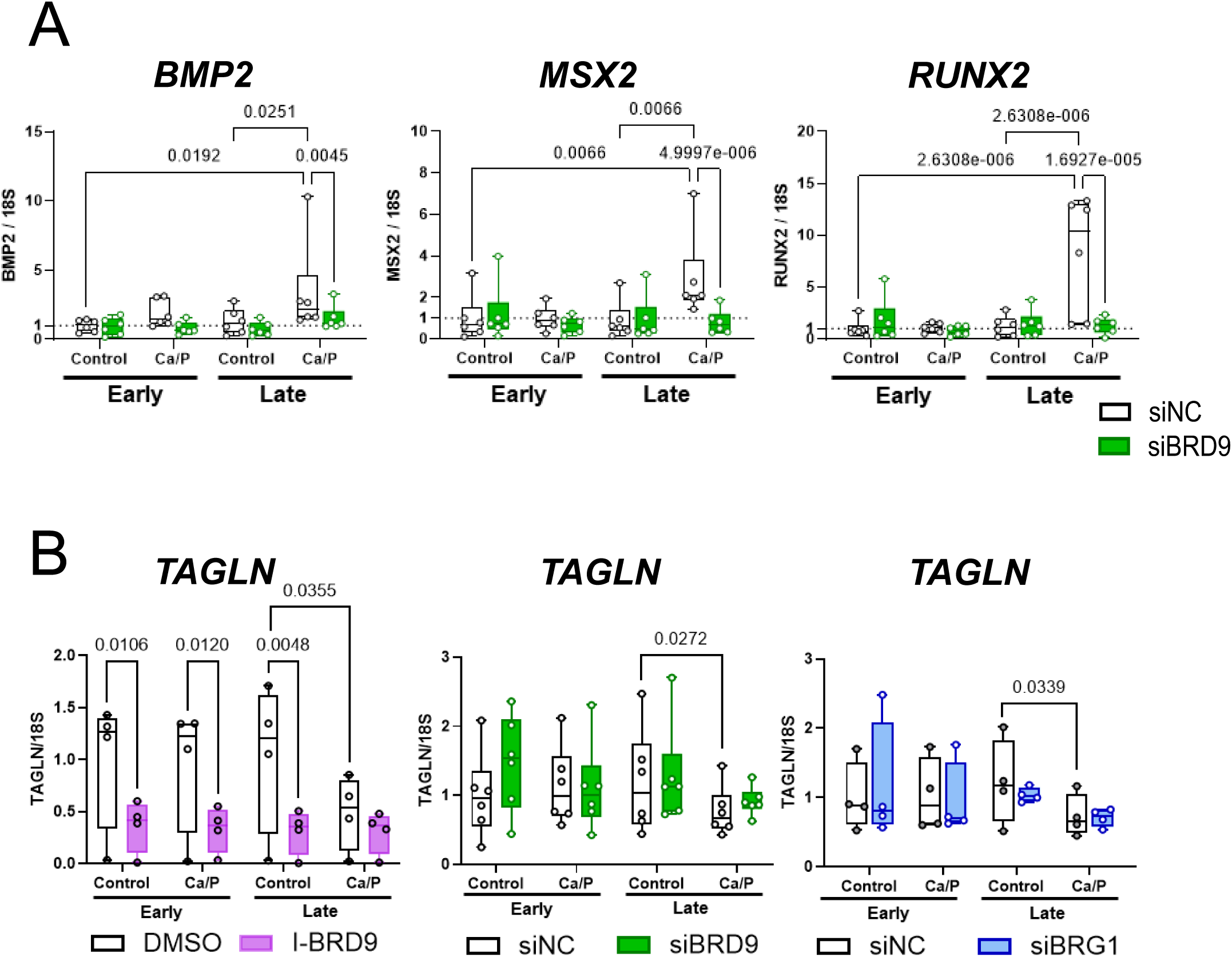
(A) Depletion of Brd9 by RNAi treatment repressed *BMP2*, *MSX2*, and *RUNX2* expression at late-stage calcification in VSMCs. VSMCs were transfected by non-targeting siRNA (siNC) or BRD9-targeting siRNA (siBRD9) under osteogenic stimuli. Samples were collected at early- or late-stage calcification. N= 6 independent experiments from 2 isolates. Significance was analyzed by Mixed-effects model test and q-values are shown. (B) Detection of *TAGLN* expression in VSMCs with inhibition of Brd9 and Brg1 under osteogenic stimuli. Samples were collected at the early- or late-stage of calcification. N= 4 independent experiments from 2 isolates for experiments with I-BRD9 treatment and loss of Brg1 by RNAi. N= 6 independent experiments from 2 isolates for experiment with depletion of Brd9 by RNAi. Significance was analyzed by Mixed-effects model test and q-values are shown.

**Supplementary Figure 3:**
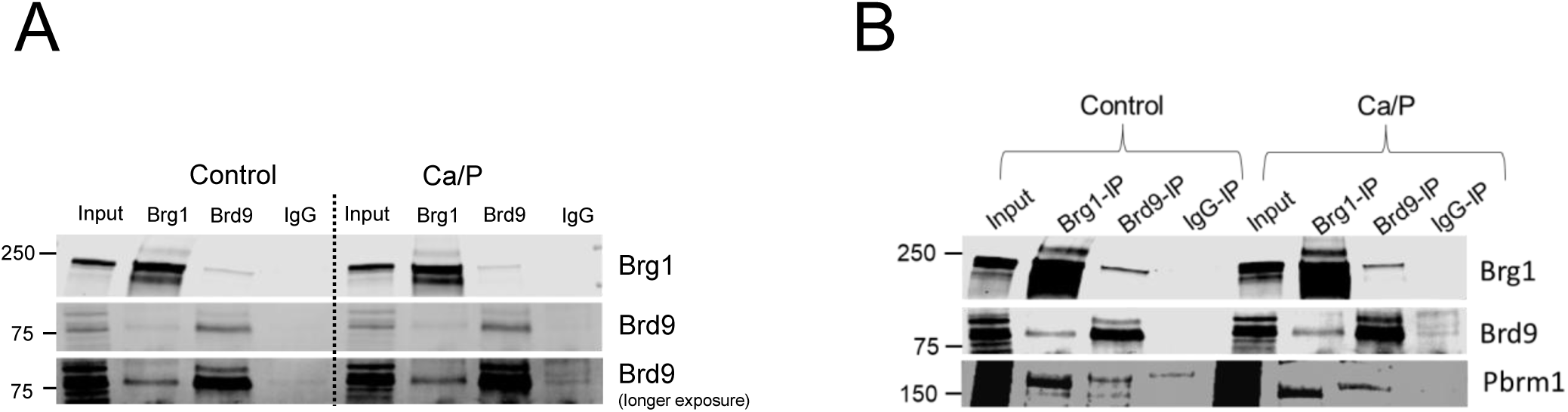
**(A)** The Brg1-Brd9 interaction was confirmed by immunoprecipitation (IP) followed by western blotting in VSMCs in the presence or absence of osteogenic stimuli. Whole cell lysates were made from VSMCs cultured in control or osteogenic media for 5 days. Antibodies against Brg1 or Brd9 were pre-bound with Protein A-dynabeads before incubation with 750 µg of whole cell extract. IP using Rabbit IgG was applied as a negative control. Capture of Brg1 and Brd9 was confirmed by western blotting. (B) The interaction between Brd9 and Pbrm1 was confirmed by immunoprecipitation (IP) followed by western blotting in VSMCs in the presence or absence of osteogenic stimuli. Capture of Brg1, Brd9, and Pbrm1 was confirmed by western blotting.

**Supplementary Figure 4:**
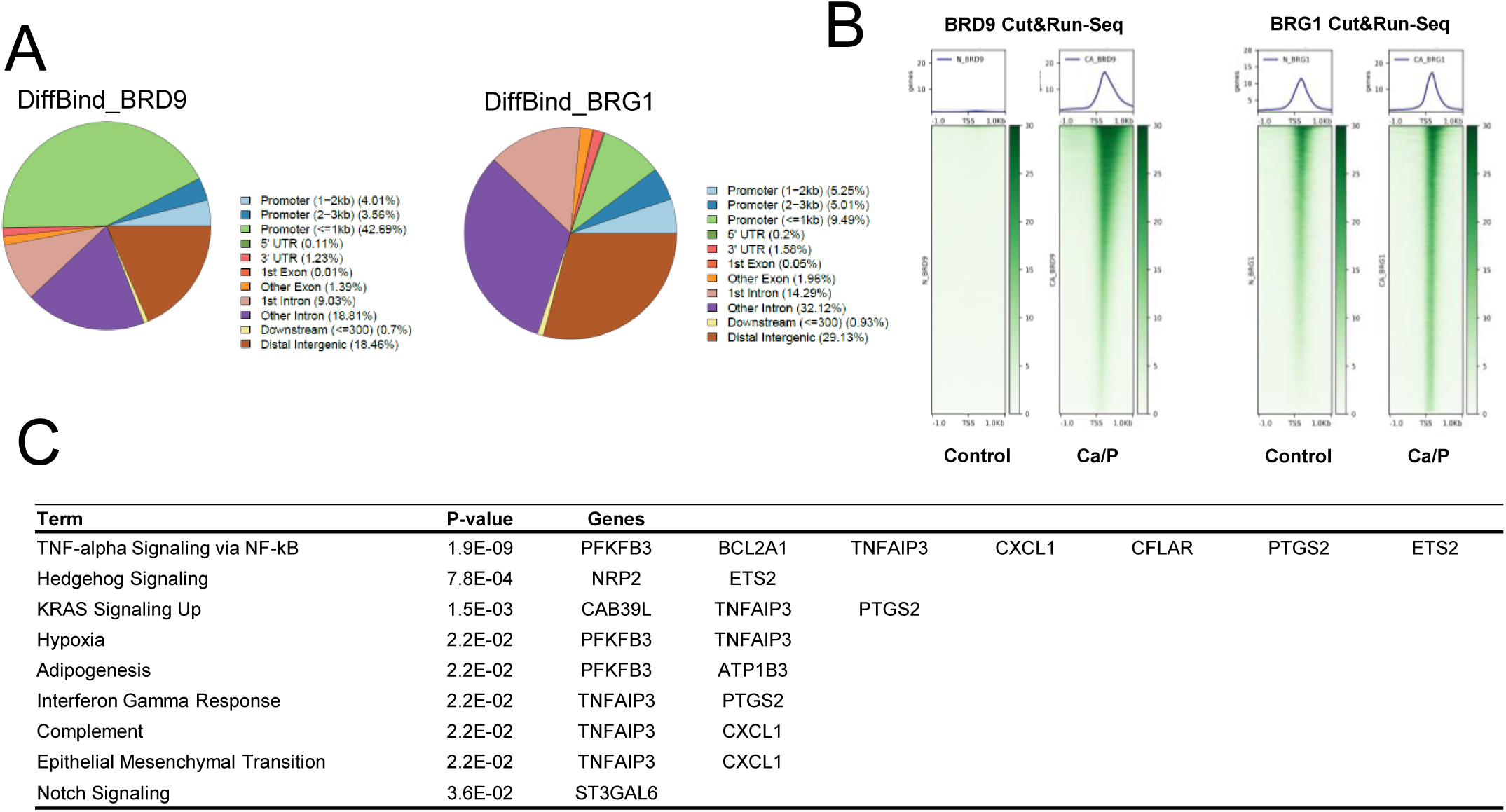
(A) Distribution of the binding regions of Brd9 (DiffBind_BRD9) and Brg1 (DiffBind_BRG1) in pre-osteogenic VSMCs. (B) Cut&Run-Seq profiles of Brd9 and Brg1 aligned at Transcription Start Sites (TSS) in control VSMCs or VSMCs in response to osteogenic stimuli (Ca/P). (C) The detailed ncBAF-dependent gene list involved in individual pathways which were activated in pre-osteogenic VSMCs.

**Supplementary Figure 5.**
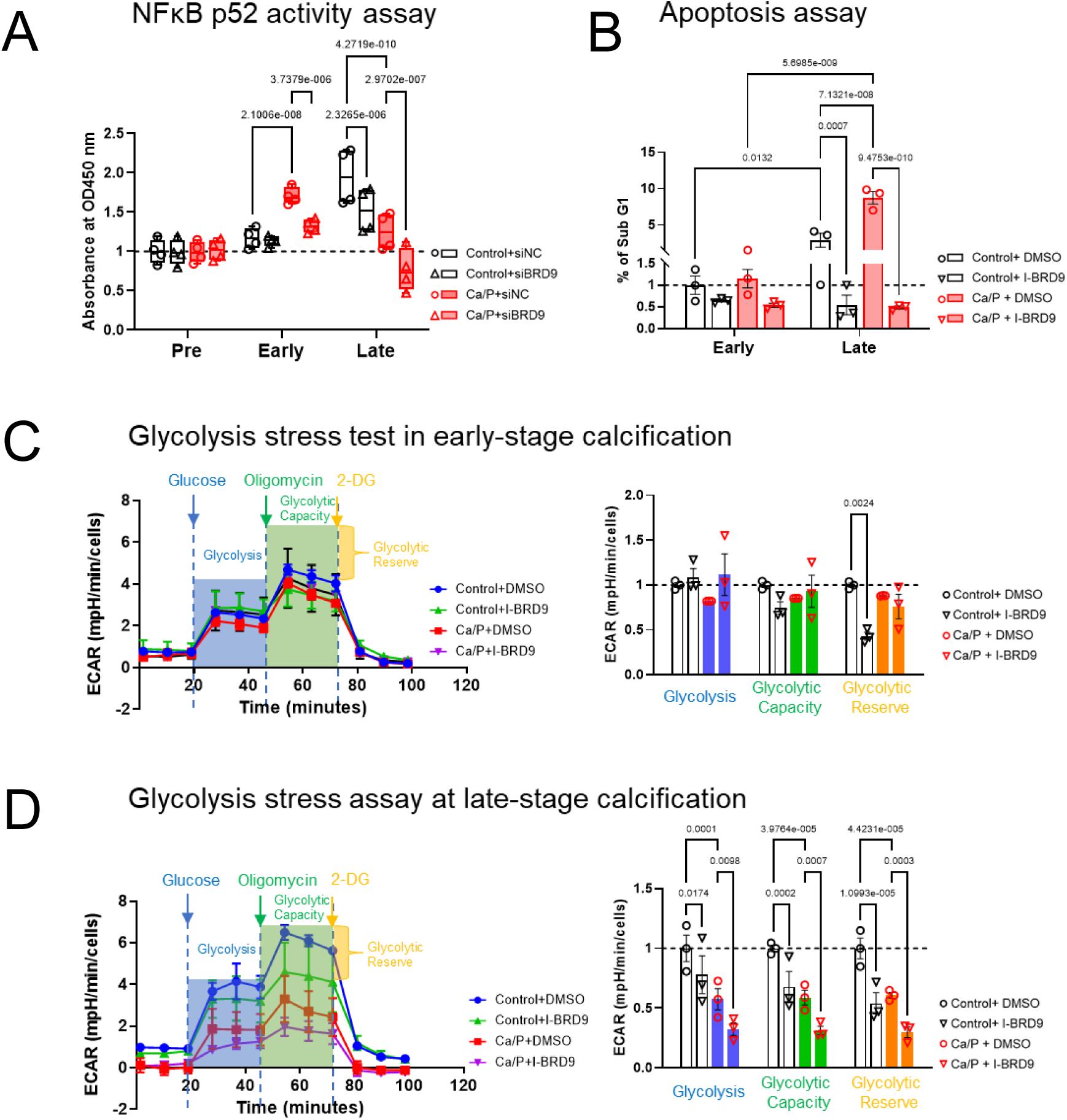
Cellular pathways activated in pre-osteogenic VSMCs were downregulated by inhibiting Brd9. (A) Quantification of NFkB p52 activation in nuclear extracts of VSMCs during the induction of osteogenic differentiation. VSMCs were treated with non-targeting siRNA (siNC) or BRD9 targeting siRNA (siBRD9) and cultured in control or osteogenic media. Cells were collected at individual stages of calcification progression followed by nuclear protein extraction. N= 4 independent experiments from 2 isolates. Significance was analyzed by Mixed-effects model test and q-values are shown. (B) The percentage of sub-G1 population which is representative of apoptotic cells was measured by DAPI staining in VSMCs cultured in control or osteogenic media in the presence or absence of Brd9 inhibitor. N= 3 independent experiments from 2 isolates. Significance was analyzed by Mixed-effects model test and q-values are shown. (C) Glycolytic function in VSMCs at early-stage calcification was measured by Seahorse glycolysis stress assay. VSMCs cultured in control or osteogenic media in the presence or absence of Brd9 inhibitor and the extracellular acidification rate (ECAR) in individual condition was measured by Seahorse XF following glycolysis stress assay. The ECAR was normalized by total cell counts. N= 3 independent experiments from 2 isolates. Significance was analyzed by Mixed-effects model test and q-values are shown. (D) Glycolytic function in VSMCs at late-stage calcification was measured by Seahorse glycolysis stress assay. VSMCs cultured in control or osteogenic media in the presence or absence of Brd9 inhibitor and the extracellular acidification rate (ECAR) in individual condition was measured by Seahorse XF following glycolysis stress assay. The ECAR was normalized by total cell counts. N= 3 independent experiments from 2 isolates. Significance was analyzed by Mixed-effects model test and q-values are shown.

**Supplementary Figure 6:**
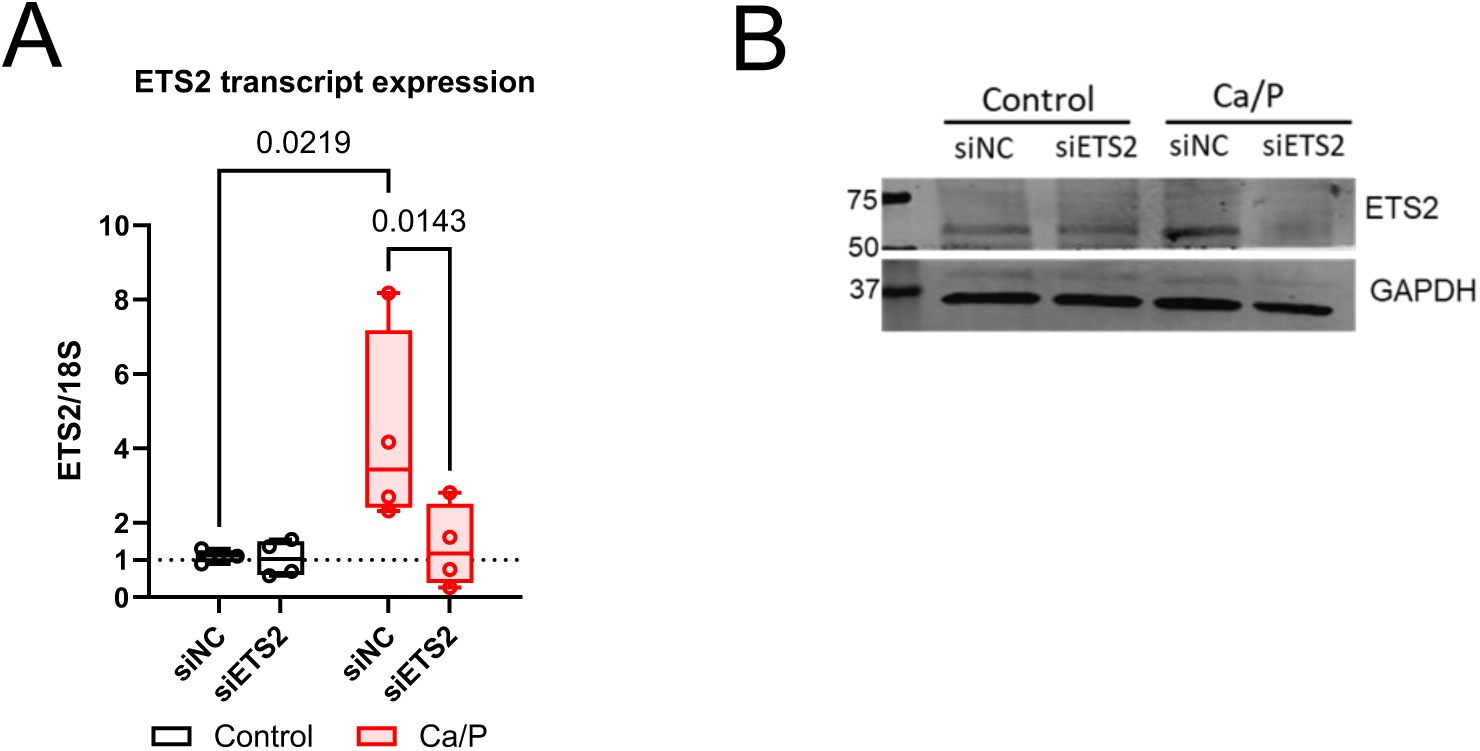
(A) Confirmation of *ETS2* knock-down efficiency by QPCR. VSMCs were transfected by non-targeting siRNA (siNC) and siRNA targeting *ETS2* (siETS2) and cultured in the presence or absence of osteogenic stimuli (Ca/P). Samples were collected at late-stage calcification and the expression of *ETS2* were detected by QPCR. N= 4 independent experiments from 2 isolates. Significance was analyzed by Mixed-effects model test and q-values are shown. (B) Western blotting showed the depletion of Ets2 by siRNA treatment under osteogenic stimuli.

**Supplementary Figure 7:**
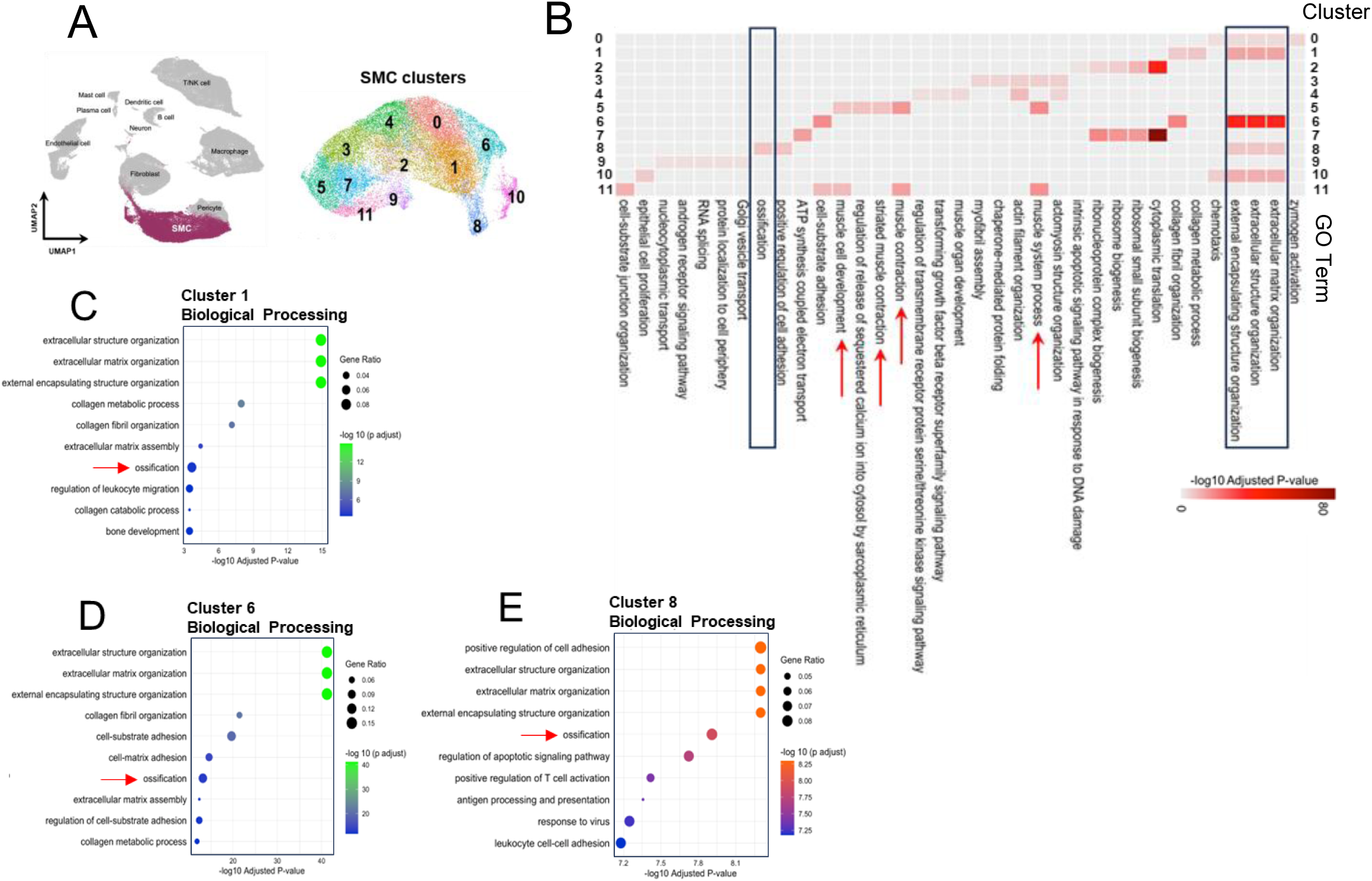
(A) Two-dimensional uniform manifold approximation and projection (UMAP) plot shows SMC clusters according to the differentially expressed genes (DEGs) in each cluster. (B) A heatmap presents the overrepresented functions of SMC clusters by gene ontology analysis of the DEGs in each cluster. (C) Biological Processing analysis of the DEGs in cluster 1 showed associations with extracellular matrix organization, ossification and bone development which are included in the top 10 scoring terms. (D) Biological Processing analysis of the DEGs in cluster 6 showed associations with extracellular matrix organization and ossification which are included in the top 10 scoring terms. (E) Biological Processing analysis of the DEGs in cluster 8 showed associations with extracellular matrix organization, ossification and T cell activation which are included in the top 10 scoring terms.

**Supplementary Figure 8.**
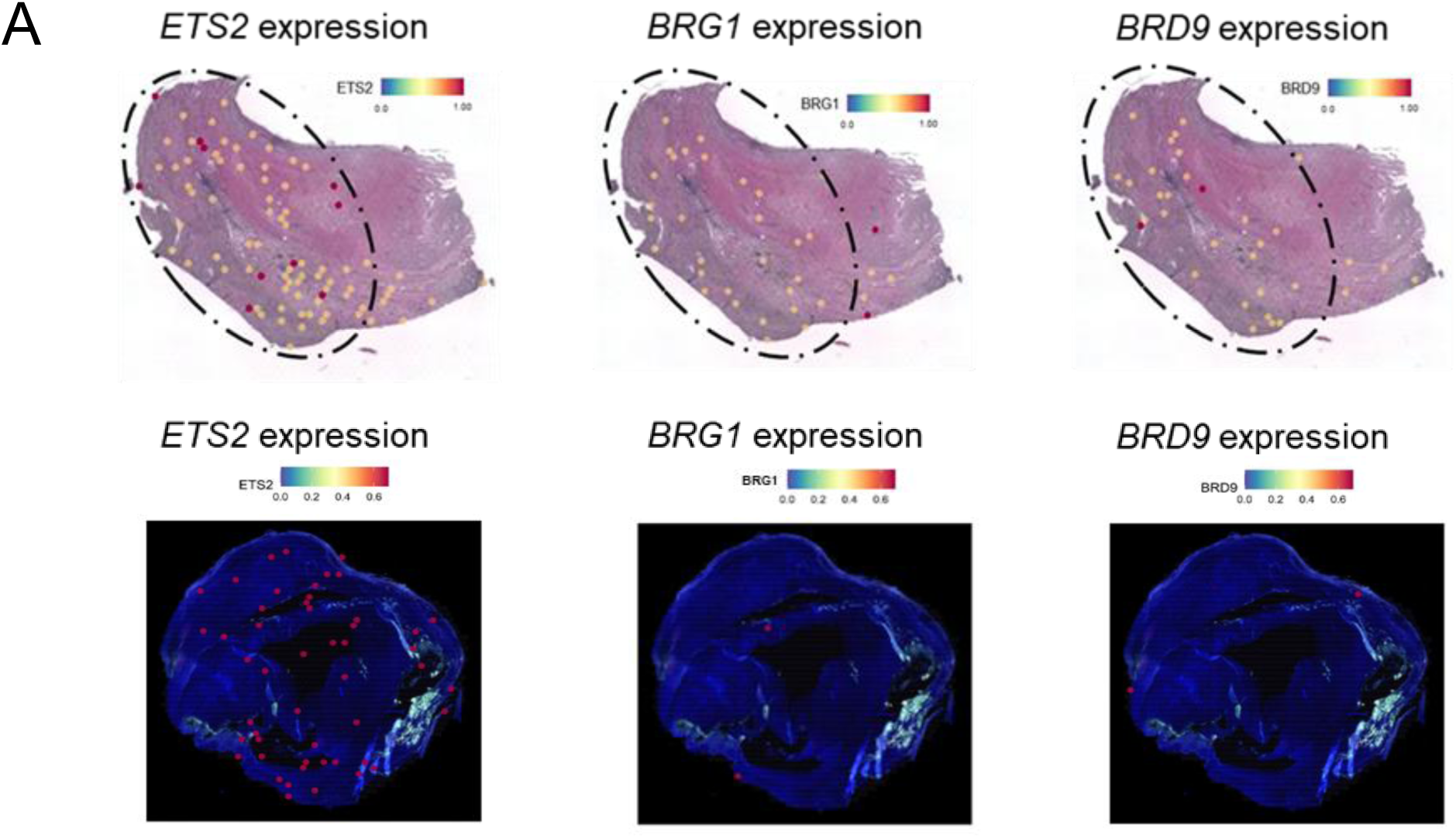
(A) Spatial expression patterns of *ETS2*, *BRG1*, and *BRD9* across the carotid plaque section Spatial #1 and #2.

