## Supplementary Table 1 for "An ncBAF-ETS2 Chromatin-Remodelling Axis Drives Vascular Smooth Muscle Cell Osteogenic Reprogramming in Vascular Calcification"

**Supplementary Table 1: List of aortic SMC isolates used in this study**

| Pt | Age (years) | Gender | Status | Vessel Type |
| --- | --- | --- | --- | --- |
| 1 | 20 | M | Normal | Aorta |
| 2 | 35 | F | Normal | Aorta |
| 3 | 38 | F | Normal | Aorta |
