## Supplementary Table 2 for "An ncBAF-ETS2 Chromatin-Remodelling Axis Drives Vascular Smooth Muscle Cell Osteogenic Reprogramming in Vascular Calcification"

**Supplementary Table 2: List of aorta specimens used in this study**

| Pt | Age (years) | Gender | Status | Vessel Type |
| --- | --- | --- | --- | --- |
| 1 | 17 | M | Normal | Aorta |
| 2 | 18 | F | Normal | Aorta |
| 3 | 19 | M | Normal | Aorta |
| 4 | 22 | M | Normal | Aorta |
| 5 | 25 | F | Normal | Aorta |
| 6 | 28 | F | Calcified | Aorta |
| 7 | 34 | M | Normal | Aorta |
| 8 | 44 | F | Calcified | Aorta |
| 9 | 45 | F | Calcified | Aorta |
| 10 | 57 | F | Calcified | Aorta |
| 11 | 63 | F | Calcified | Aorta |
| 12 | 64 | F | Calcified | Aorta |
| 13 | 67 | M | Calcified | Aorta |
| 14 | 70 | M | Calcified | Aorta |
| 15 | 76 | M | Normal | Aorta |
| 16 | 89 | F | Calcified | Aorta |
