## Supplementary Table 3 for "An ncBAF-ETS2 Chromatin-Remodelling Axis Drives Vascular Smooth Muscle Cell Osteogenic Reprogramming in Vascular Calcification"

**Supplementary Table 3. Primers**

| **RT-qPCR primers** | |
| --- | --- |
| 18S ribosomal RNA | QT00199367 /Hs_RRN18S_1_SG QuantiTect Primer Assay |
| RUNX2 | QT00020517 Hs_RUNX2_1_SG QuantiTect Primer Assay |
| IL6 | QT00083720 Hs_IL6_1_SG QuantiTect Primer Assay |
| TAGLN | QT00072247 Hs_TAGLN_1_SG QuantiTect Primer Assay |
| BRG1 | QT00046578 Hs_SMARCA4_1_SG QuantiTect Primer Assay |
| BMP2 | QT00012544 Hs_BMP2_1_SG QuantiTect Primer Assay |
| BRD9 forward | 5' GCC ACG ACT CCA GTT AC 3' |
| BRD9 reverse | 5' GAT GCT TCT CCT TCT CGG 3' |
| MSX2 forward | 5’ AAATTCAGAAGATGGAGCGGCGTG 3’ |
| MSX2 reverse | 5’ CGGCTTCCGATTGGTCTTGTGTTT 3’ |
| ETS2 forward | 5' CTC CGT TCC TCA TTG GAT 3' |
| ETS2 reverse | 5' CGC CTT TGG GGT AAT TCT 3' |
| TNFAIP3 forward | 5' CAC AAT GGC TGA ACA AGT C 3' |
| TNFAIP3 reverse | 5' AAA TGT CTT CTG GAG TTC TCT 3' |
| CXCL1 forward | 5' CGA AGT CAT AGC CAC ACT C 3' |
| CXCL1 reverse | 5' TTG TCA CTG TTC AGC ATC T 3' |
| PTGS2 forward | 5' CAA GTC CCT GAG CAT CTA C 3' |
| PTGS2 reverse | 5' ATA CTC TGT TGT GTT CCC G 3' |
| BCL2A1 forward | 5' AGT CAT GCT TGG ACA ATG TTA 3' |
| BCL2A1 reverse | 5' TGC CGT CTT CAA ACT CCT 3' |
| PFKFB3 forward | 5' CAG GAG AAT GTG CTG GTC 3' |
| PFKFB3 reverse | 5' GGC ATT TCA GGT AGG GC 3' |
| **ChIP-qPCR primers** | |
| BMP2 promoter_For | 5'GAG CAG GGA GTG GAG GC 3' |
| BMP2 promoter_Rev | 5' GCA GGG GTG TGG ACG GC 3' |
| MSX2 promoter_For | 5' CTG ACT GCT CCT GTA ATT AAC TC 3' |
| MSX2 promoter_Rev | 5 'CTG ATT GGC TCT TCC CGA 3' |
| RUNX2 promoter_For | 5' AAG AGA GAG AGA GAA AGA GCA A 3' |
| RUNX2 promoter_Rev | 5' TAT TAC TGG AGA GGC AGA ATC AT 3' |
| ETS2 promoter_For | 5' GGC TTG ATC GTA GTT CAC C 3' |
| ETS2 promoter_Rev | 5' GGC TTA TGC CTG TAA CCC 3' |
